# Spatial and temporal inhibition of FGFR2b ligands reveals continuous requirements and novel targets in mouse inner ear morphogenesis

**DOI:** 10.1101/430876

**Authors:** Lisa D. Urness, Xiaofen Wang, Huy Doan, Nathan Shumway, C. Albert Noyes, Edgar Gutierrez-Magana, Ree Lu, Suzanne L. Mansour

## Abstract

Morphogenesis of the inner ear epithelium requires coordinated deployment of several signaling pathways and disruptions cause abnormalities of hearing and/or balance. The FGFR2b ligands, FGF3 and FGF10, are expressed throughout otic development and are required individually for normal morphogenesis, but their prior and redundant roles in otic placode induction complicates investigation of subsequent combinatorial functions in morphogenesis. To interrogate these roles and identify new effectors of FGF3 and FGF10 signaling at the earliest stages of otic morphogenesis, we used conditional gene ablation after otic placode induction and temporal inhibition of signaling with a secreted, dominant-negative FGFR2b ectodomain. We show that both ligands are required continuously after otocyst formation for maintenance of the otic ganglion and patterning and proliferation of the epithelium, leading to normal morphogenesis of both the cochlear and vestibular domains. Furthermore, the first genomewide identification of proximal targets of FGFR2b signaling in the early otocyst reveals novel candidate genes for inner ear development and function.

## INTRODUCTION

The membranous labyrinth of the mammalian inner ear is among the most complex examples of organ morphogenesis. An unremarkable patch of cranial ectoderm is transformed into a structurally intricate sensory apparatus with two functionally distinct compartments: the ventral cochlea and the dorsal vestibular system, responsible for the perception of sound and acceleration, respectively. Within these compartments an exquisitely patterned array of sensory, non-sensory and supporting cell types are poised to transduce auditory and vestibular stimuli through sensory ganglia to the brain. Proper morphogenesis of the labyrinth is essential for normal auditory and vestibular function as indicated by imaging studies of hearing loss patients (Kimura et al., 2018; Sennaroglu and Bajin, 2017). In light of the advent of cochlear implantation to treat hearing loss in cases of inner ear malformation (Isaiah et al., 2017), elucidating signals governing otic morphogenesis and appreciation of the spectrum of labyrinthine morphogenetic defects are necessary to advance treatment.

Amniote inner ear development initiates during neurulation when hindbrain proximal ectoderm is induced to thicken, forming the otic placode, the source of both the otic epithelium and the neurons of its sensory ganglia. Next, the placode invaginates, forming a cup that deepens and delaminates neuroblasts, before pinching off from the overlying ectoderm to form a spherical vesicle, the otocyst, that at embryonic day (E)9.5 in mouse is already patterned along the three anatomical axes. Otocyst morphogenesis initiates with dorsomedial budding to form the endolymphatic duct and sac (EDS) anlagen. Vestibular structures initiate by sequential dorsal and lateral evaginations of the epithelium to form vertical and horizontal pouches, which are further sculpted by epithelial fusion and resorption, generating the three orthogonal semicircular canals. The utricle and saccule form from anterior/central bulges, and the cochlear duct (CD) emerges as a ventral outgrowth. In mice, it undergoes progressive ventral extension and coiling, reaching 1.75 turns by E15.5, when gross morphogenesis is largely complete (Morsli et al., 1998; Sajan et al., 2007). Cell type differentiation and functional maturation however, continue until well after birth (reviewed in Alsina and Whitfield, 2017; Basch et al., 2016; Whitfield, 2015; Wu and Kelley, 2012).

Signals regulating otic placode induction and early otocyst patterning emanate from surrounding tissues and are understood in some detail (Ladher, 2017), but by otocyst stages, intrinsic signals are also produced and their roles in driving region-specific morphogenesis are less well understood. Extrinsic signals regulating dorsal morphogenesis include hindbrain WNTs and BMPs. SHH secreted from the ventral hindbrain and notochord initiates ventral morphogenesis. Crosstalk between these signals involves regulation of key region-specific transcription factors (Ohta and Schoenwolf, 2018). FGF signaling also plays critical roles in otic development and functions at multiple stages. A cascade of FGFs from endoderm, mesoderm and hindbrain is required for otic placode induction (Alvarez et al., 2003; Ladher et al., 2005; Wright and Mansour, 2003a; Zelarayan et al., 2007). In particular, *Fgf3* and *Fgf10*, encoding ligands that signal through the same FGF receptor isoforms (FGFR1b and FGFR2b; Zhang et al., 2006), are required redundantly for otic placode induction, such that germline double null mutants (F3KO;F10KO) have no inner ear (Alvarez et al., 2003; Wright and Mansour, 2003a). Applications of FGFs and FGFR inhibitors to chick embryos revealed profound influences of FGFs on otic morphogenesis (Chang et al., 2004) and studies of individual mouse mutants revealed roles for *Fgf3* and *Fgf10* in morphogenesis. Mice lacking *Fgf3* (F3KO) fail with variable penetrance and expressivity to form an EDS, and consequently have variable morphogenesis and dysfunction of both the cochlea and vestibule (Hatch et al., 2007; Mansour et al., 1993). Mice lacking *Fgf10* (F10KO) fail to form posterior semicircular canals (PSCCs), and have milder deformations of the anterior and lateral canals. *Fgf10* heterozygotes also exhibit PSCC reductions or agenesis (Pauley et al., 2003; Urness et al., 2015). The CD is also affected in *Fgf10* null mutants, being shorter and narrower than that of heterozygous or wild-type mice due to loss of non-contiguous non-sensory domains (Urness et al., 2015). *Fgfr2b* null mutants form otocysts (Pirvola et al., 2000), however, they develop with severe cochlear and vestibular dysmorphology, suggesting that *Fgf3* and *Fgf10* could have additional and combinatorial roles during morphogenesis.

Here, we define the dynamic expression of *Fgf3* and *Fgf10* in the developing mouse otic epithelium and ganglion, and interrogate their functions after otic placode induction. We employ two complementary genetic strategies: otic placode lineage-restricted gene ablation and timed induction of a soluble dominant-negative FGFR2b ectodomain that acts rapidly as an extracellular ligand trap to block signaling. Together, our data show that *Fgf3* and *Fgf10* are not required in the placode lineage for otocyst formation, but are required subsequently for otocyst patterning, neuroblast maintenance, epithelial proliferation and both vestibular and cochlear morphogenesis. Furthermore, the first differential RNA-Seq analyses of otocysts revealed FGFR2b signaling targets that define novel candidates for genes involved in otic morphogenesis and function.

## RESULTS

### *Fgf3* and *Fgf10* are expressed dynamically during otocyst and ganglion formation, and cochlear morphogenesis

To determine *Fgf3* and *Fgf10* expression during otocyst formation and cochlear morphogenesis, we used RNA in situ hybridization (ISH). Before E9.0, both genes were exclusively periotic (data not shown). Consistent with previous studies (Schimmang, 2007; Wright and Mansour, 2003b), *Fgf3* and *Fgf10* transcripts were non-overlapping at the otic cup stage, with *Fgf3* expressed in hindbrain adjacent to the cup (Fig. 1A), and *Fgf10* expressed in the cup itself, exclusive of the dorsal and lateral-most regions (Fig. 1B). Once the otocyst closed, *Fgf3* was diminished in the hindbrain and was first seen in the ventrolateral otocyst and in the forming otic ganglion (Fig. 1C), whereas *Fgf10* was expressed in the ventral and medial otocyst (Fig. 1D). By E10.25-E11.25, *Fgf3* was confined to the ventrolateral otocyst (Figs. 1E,G). At this stage, *Fgf10* overlapped with and was more extensively expressed in the ventral otocyst than was *Fgf3*, and also began to be expressed strongly in the ganglion (Figs. 1F,H).

**Figure 1.**
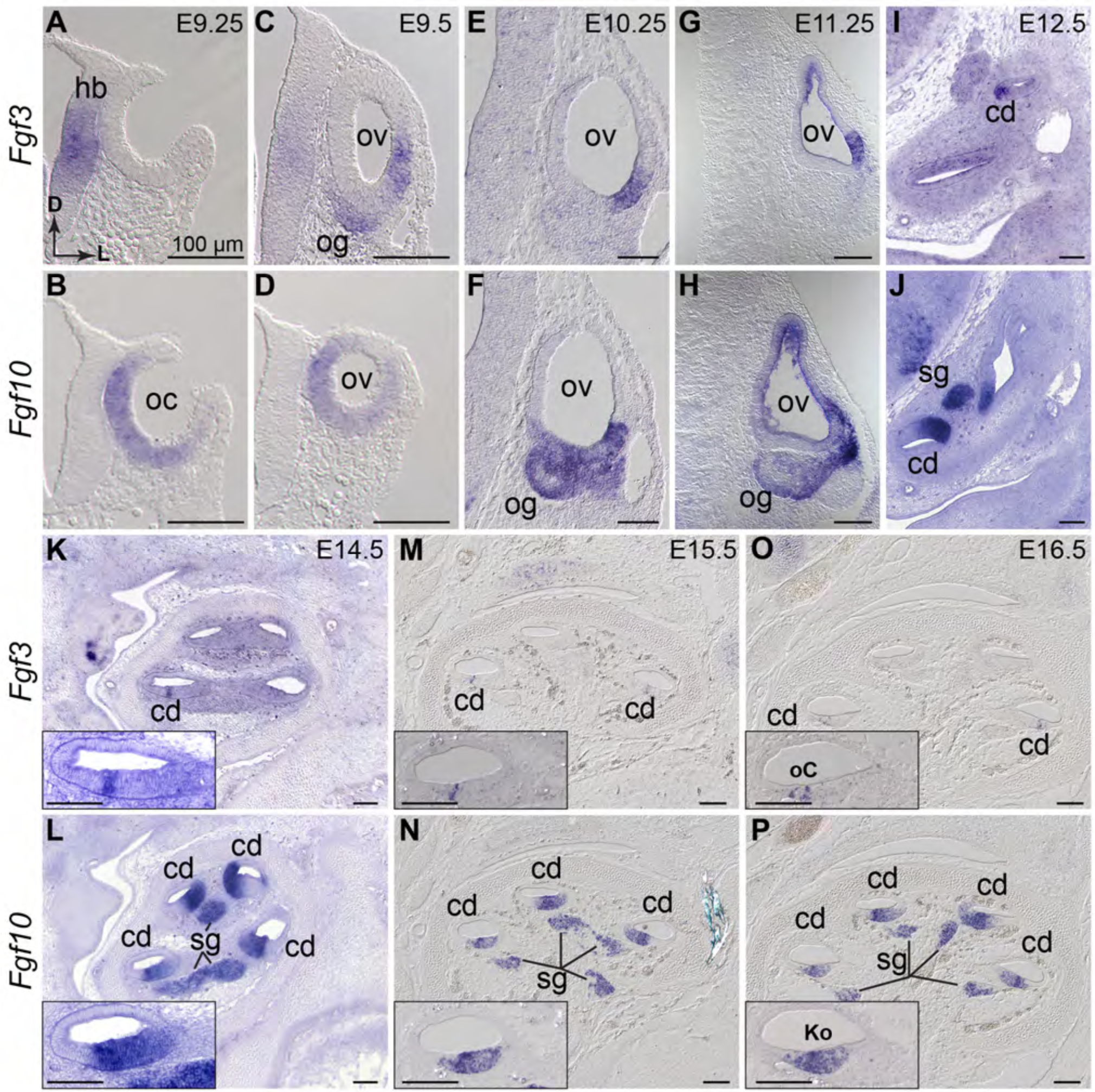
*Fgf3* and *Fgf10* are expressed throughout otocyst formation and cochlear morphogenesis. ISH of *Fgf3* and *Fgf10* probes to wild type otic cup (A,B), otic vesicle (C-H), and cochlear duct (I-P). Probes indicated to the left and developmental ages at the top right of each paired column. (A-J) transverse orientation, directional arrows in A apply to all; (K-P) sagittal orientation with cochlear apex at the top. Insets show the basal-most cochlear turn. Scale bars indicate 100 μm. Abbreviations: cd, cochlear duct; D, dorsal; hb, hindbrain; Ko, Kölliker’s organ; L, lateral; oC, organ of Corti; oc, otic cup; og, otic ganglion; ov, otic vesicle; sg, spiral ganglion.

We confirmed overlap of *Fgf3* and *Fgf10* in the developing vestibular sensory tissues, with *Fgf10* expression much stronger than *Fgf3* (data not shown; see Pauley et al., 2003; Pirvola et al., 2000). Therefore, we focused on the developing CD. From E12.5-E16.5, we found *Fgf3* in a progressively limited portion of the CD that appeared by E16.5 to flank the developing sensory organ of Corti (Figs. 1I,K,M,O). *Fgf10* continued expression in a broader cochlear epithelial domain than *Fgf3*, resolving to Kölliker’s organ by E16.5, and was maintained at high levels in the cochlear ganglion (Figs. 1J,L,N,P). These observations suggested that after otic induction, *Fgf3* and *Fgf10* could have combinatorial roles in morphogenesis and ganglion development.

### Epithelial *Fgf3* and *Fgf10* are not required for otocyst formation, but both are required for vestibular and cochlear morphogenesis

Since F3KO;F10KO embryos lack inner ears (Alvarez et al., 2003; Wright and Mansour, 2003a), we took a conditional approach to disrupt these genes individually and combinatorially after otic placode induction using *Tg(Pax2-Cre)*, which is active in the otic placode lineage starting at E8.5 (Fig. 2A; Ohyama and Groves, 2004). As this lineage comprises both epithelium and ganglion, Tg(Pax2-Cre) recombines in both tissues (Figs. 2B,C). In contrast to the variable otic phenotypes observed in F3KO mutants (Hatch et al., 2007; Mansour et al., 1993), deletion of *Fgf3* alone in the *Pax2-Cre* lineage (F3cKO) had no effect on otic morphogenesis at E15.5 (Figs. 2D,D’) or on CD histology at E18.5 (Figs. 2E,E’). Indeed, F3cKO animals survived in the expected numbers and had normal auditory thresholds and motor behavior (data not shown). In contrast, F10cKO ears showed both vestibular and cochlear abnormalities, including reduction or loss of the PSCC and variable shortening and narrowing of the CD (Figs. 2F,F’), reflecting loss of Reissner’s membrane (Figs. 2G,G’). These abnormalities were similar to those of F10KO ears (Urness et al., 2015) and, indeed, immunostaining of E18.5 F10cKO CDs was similar to F10KO CDs (data not shown). Notably, *Fgf10^-/c^* Cre-negative ears had mild PSCC shortening (Fig. 2F), but CD defects appeared only in F10cKO ears (Fig. 2F’).

**Figure 2.**
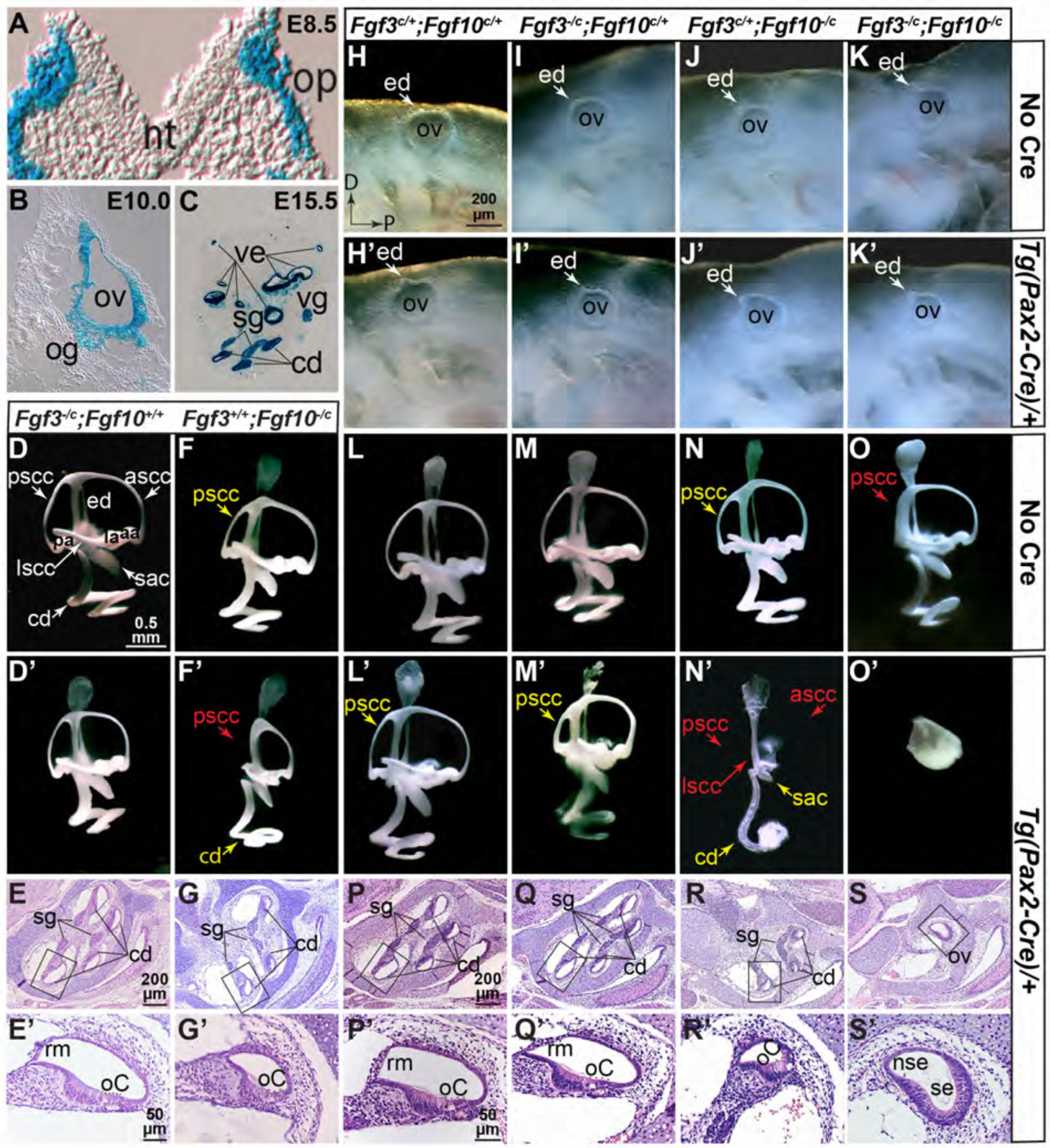
*Fgf3* and *Fgf10* are not required in the *Pax2-Cre* lineage for otocyst formation, but both are required subsequently for vestibular and cochlear morphogenesis. *Pax2-Cre* otic lineage (blue) at E8.5 (A), E10.0 (B) and E15.5 (C). E9.5 left lateral otocysts (H-K’). E15.5 paint filled right lateral inner ears (D-F’,L-O’) and E18.5 H&E-stained sagittal cochlear sections (E,G,P-S). Enlargements of boxed regions shown in E’,G’,P’-S’. *Fgf* genotypes shown above each column and *Cre* allele status indicated to the right of each row. Text in yellow indicates apparent length reductions, text in red indicates expected positions of missing structures. Scale bar in H applies to H-K’, in D to D-F’,L’-O’, in E, to E,G, in P to P-S, in E’ to E’,G’ and in P’ to P’-S’. Abbreviations not used previously: aa, anterior ampula; ascc, anterior semicircular canal; ed, endolymphatic duct; la, lateral ampula; lscc, lateral semicircular canal; nt, neural tube; nse, nonsensory epithelium; pa, posterior ampula; pscc, posterior semicircular canal; rm, Reissner’s membrane; sac, saccule; se, sensory-like epithelium, vg, vestibular ganglion; ve, vestibular epithelium.

Next, we evaluated morphogenesis after varying *Fgf3* and *Fgf10* allele dosage. At E9.5, embryos of all genotypes had otocysts starting to develop an EDS (Figs. 2H,H’-K,K’), showing that, as expected from the timing of CRE activity onset, otic placode induction occurred normally, even in the absence of both *Fgfs* (Fig. 2K’). In contrast, when analyzed at E15.5 by paintfilling, distinct defects in epithelial morphology were apparent in many different genotypes (Figs. 2L,L’-O,O’). Even Cre-negative ears (Figs. 2L-O) showed vestibular defects when heterozygous for the *Fgf10* null allele (Figs. 2N,O). Not surprisingly, all Cre-positive ears had reduced or absent PSCCs, as *Fgf10* is at least heterozygous in those cases (Figs. 2L’-O’). These ears formed an allelic series of increasing severity, with F3cHet;F10cHet ears showing only mild PSCC reductions (Fig. 2L’). This was exacerbated in F3cKO;F10cHet ears (Fig. 2M’). However, F3cHet/F10cKO ears lost virtually the entire SCC system, showed reductions in the saccule and utricle, and had a more extreme shortening and narrowing of the CD (Fig. 2N’) than did F10cKO ears (Fig. 2F’). Only the EDS appeared normal. Preliminary analyses of E18.5 F3cHet/F10cKO ears did not reveal any exacerbation of changes in cochlear marker genes analyzed previously in F10KO ears (data not shown, see Urness et al., 2015). Strikingly, conditional disruption of both *Fgf3* and *Fgf10* blocked both vestibular and cochlear development, leaving only a small spherical vesicle (Fig. 2O’). Histologic sections of E18.5 F3cHet;F10cHet and F3cKO;F10cHet CDs (Figs. 2P,P’,Q,Q’) were indistinguishable from those of F3cKO CDs (Fig. 2E,E’), whereas F3cHet;F10cKO CDs were very narrow and lacked Reissner’s membrane (Fig. 2R,R’), similar to those of F10cKO (Fig. 2G,G’) and F10KO CDs (Urness et al., 2015). Whether the extreme narrowing of the CD in F3cHet;F10cKO ears is consistently more severe than in other types of *Fgf10* mutants is unknown. The F3cKO;F10cKO “ear” had an epithelium comprising a thin, non-sensory region and a thickened vestibular-like sensory region. Most notably, these mutants showed no evidence of cochlear or vestibular neurons (Figs. 2S,S’). These data show that both *Fgf3* and *Fgf10* are required in the *Pax2-Cre* lineage, not only for vestibular morphogenesis, but also for cochlear morphogenesis and otic gangliogenesis, with the role of *Fgf3* being revealed only in the absence of *Fgf10*.

### *Fgf3* and *Fgf10* are not required in the *Pax2-Cre* lineage for early otocyst proliferation, but are required for otocyst patterning and maintenance of otic neuroblasts

Since F3cKO;F10cKO embryos ultimately develop very small otic vesicles, and this was first apparent at E10.5-E11.5, we quantified mitotic cells in E10.5 otocyst sections by calculating the number of phosphohistone H3 (pHH3)-positive cells per otic epithelial area. Somewhat surprisingly, the difference between control and F3cKO;F10cKO vesicles was not significant (Fig. S1A-C), perhaps as a consequence of *Fgf3* maintained in the hindbrain through at least E9.25 (Fig. 1A). Indeed, in the absence of *Fgf10, Fgf3* is required in the *Sox1^Cre^* lineage (hindbrain) to form a normally sized otocyst (Fig. S1D-G’), but this source of *Fgf3* is not affected by *Pax2-Cre* (Fig. 2A).

To assess otocyst patterning in conditional mutants, we conducted whole-mount ISH analyses of E9.5-E11.5 samples using probes to detect regionally expressed genes that are known targets of FGF3 and/or FGF10 signaling and/or are required for morphogenesis. To manage the number of samples analyzed, we omitted single conditional mutants and used only a single *Cre*-negative genotype *(Fgf3^-/c^;Fgf10^-/c^)*. Stained embryos were sectioned through the otocyst or viewed as whole mounts. At E9.5 most genes tested, as exemplified by *Sox9*, were unaffected even in F3cKO;F10cKO ears (Figs. 3A_1_-A_5_). Other genes unaffected by loss both *Fgf3* and *Fgf10* alleles at this stage included *Dusp6, Spry1, Foxg1, Has2, Gbx2, Hmx3, Sox2* and *Pax2* (data not shown), all of which are lost in F3KO;F10KO ears by otic placode stages (Alvarez et al., 2003; Urness et al., 2010; Wright and Mansour, 2003a). However, the common FGF signaling target, *Etv5*, which has distinct ventromedial and dorsolateral domains in E9.5 control otocysts (Fig. 3B_1_, B_2_), showed differential localization in conditional mutants with only a single *Fgf3* or *Fgf10* allele remaining (Figs. 3B_3_,3B_4_), and expression was entirely absent from F3cKO;F10cKO ears (Fig. 3B_5_), demonstrating that epithelial FGF3/FGF10 signaling was disrupted within 24 hours of CRE activation. The only other gene affected at E9.5 was *Tbx1*, which had dorsolateral and posteroventral otocyst domains in all genotypes (Figs. 3C_1_–3C_4_) except F3cKO;F10cKO, which lost the dorsolateral domain (Fig. 3C_5_). By E10.5, the dorsolateral *Tbx1* domain was lost from both F3cHet;F10cKO and F3cKO;F10cKO otocysts (Figs. 3D_4_,D_5_).

**Figure 3.**
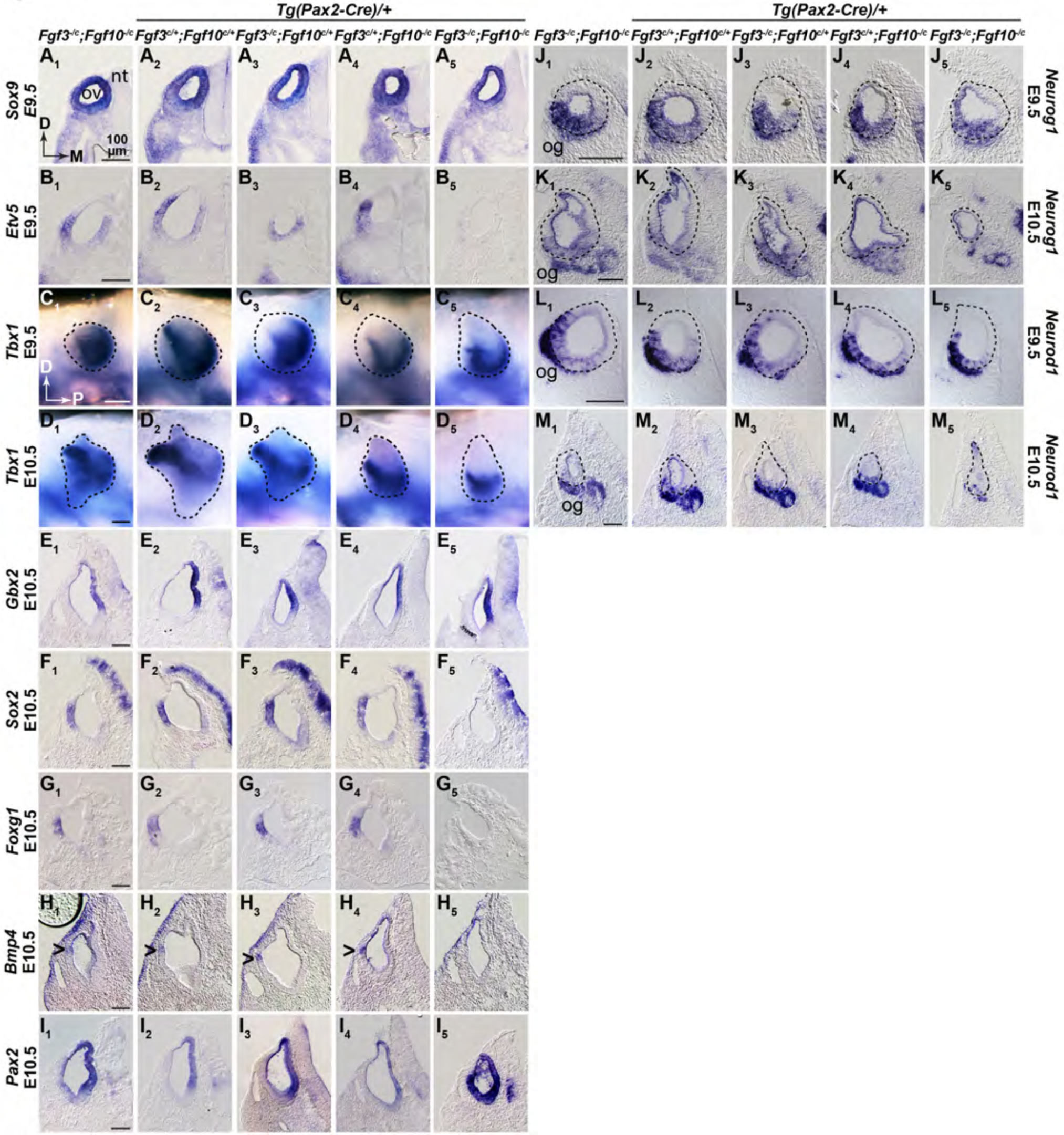
*Fgf3* and *Fgf10* are required in the *Pax2-Cre* lineage for otocyst patterning and maintenance of otic neuroblasts. Transverse sections through the ov (A,B,E-M), or whole mounts (C,D), of embryos subjected to ISH. Probes and developmental stages are indicated aside each row. Genotypes are shown atop each column. Scale bars indicate 100 μm and apply to each probe/stage group. Carets in H_1_-H_4_ indicate *Bmp4* expression. Directional arrows shown in A_1_ and C_1_ apply to all sections and whole-mounts, respectively. Abbreviations defined in Figs. 1,2.

At E10.5, expression of several other genes was still unaffected, as exemplified by *Gbx2* (Fig. 3E_1_-E_5_), which is expressed dorsomedially, is down-regulated at this stage in F3KO mutants (Hatch et al., 2007), and is required for vestibular morphogenesis (Lin et al., 2005). Other genes unaffected by loss of *Fgf3* and *Fgf10* at E10.5 included *Hmx3, Spry2, Gli1, Id1*, and *Lfng* (data not shown). In contrast, *Sox2, Foxg1* and *Bmp4*, which are primarily lateral at E10.5, were all unaffected, except in F3cKO;F10cKO ears, where expression was extinguished (Figs. 3F_5_,G_5_,H_5_). Other genes lost from E10.5 F3cKO/F10cKO otocysts included *Dusp6, Etv5, Etv4*, and *Spry2* (data not shown), all of which are known transcriptional targets of FGF signaling. Curiously, we found that *Pax2*, which at E10.5 is normally expressed medially (Fig. 3I_1_), was unchanged in all otocysts except F3cKO/F10cKO, where it was expanded (Figs. 3I_1_-I_5_).

By E11.5 the only tested genes still unaffected in F3cKO;F10cKO otocysts were *Gli1, Hmx3, Lfng* and *Id1* (data not shown), so these are unlikely to be targets of FGF3/FGF10 signaling in the otocyst. Whether the affected genes are direct or indirect targets of FGF signaling at this stage could not be determined.

To assess the otic ganglion, we assayed *Neurog1* and its target, *Neurod1*, at both E9.5 and E10.5. At E9.5, all genotypes exhibited similar expression in a ventrolateral epithelial domain and in delaminating neuroblasts (Figs. 3J_1_-J_5_,3L_1_-L_5_). In contrast, at E10.5 *Neurog1* was strongly reduced and *Neurod1* was virtually eliminated from the otic epithelium, and otic ganglion development was suppressed specifically in F3cKO/F10cKO otocysts (Figs. 3K_5_,M_5_). These data show that epithelial/ganglion expression of *Fgf3* and *Fgf10* are required for aspects of gene expression driving otic morphogenesis, particularly in lateral regions, and that they are also required for otic ganglion formation.

### Doxycycline induction of a secreted FGFR2b ectodomain phenocopies *Fgf3/Fgf10* double null mutants

FGF3 and FGF10 bind to and signal primarily through “b”-type FGF receptors (FGFR2b>FGFR1b, Zhang et al., 2006). To enable simultaneous and inducible inhibition of their signaling activity at any stage, we employed two alleles that together enable doxycycline (DOX)-inducible expression of a secreted, dominant-negative form of FGFR2b (dnFGFR2b), which serves as a ligand trap. *Rosa26^rtTA^* drives ubiquitous expression of the reverse tetracycline transactivator (Belteki et al., 2005) and *Tg(tetO-dnFgfr2b)* encodes a tetO-regulated and secreted FGFR2b ectodomain (Hokuto et al., 2003). This system is validated for temporally controlled inhibition of mammary gland, tooth, limb and lung development, which depend on various FGFR2b and FGFR1b ligands (Al Alam et al., 2015; Danopoulos et al., 2013; Parsa et al., 2010; Parsa et al., 2008).

To validate this system for inner ear studies we crossed the alleles together, fed DOX-chow to pregnant females from E5.5-E10.5 and observed gross embryonic phenotypes. Relative to *Rosa26^rtTA/+^* embryos, which appeared normal (Fig. 4A), double heterozygotes had a short curly tail, lacked limb buds and had tiny otic vesicles (Fig. 4B), phenocopying F3KO;F10KO mutants. Double heterozygotes exposed to DOX from E5.5 or E6.5 to E11.5 showed only an otic remnant (Fig. S2) and they did not exhibit mid-hindbrain phenotypes characteristic of inhibition of ligands such as FGF8 and FGF17, that signal through “c”-type FGFRs (Chi et al., 2003; Sato and Joyner, 2009; Xu et al., 2000).

**Figure 4.**
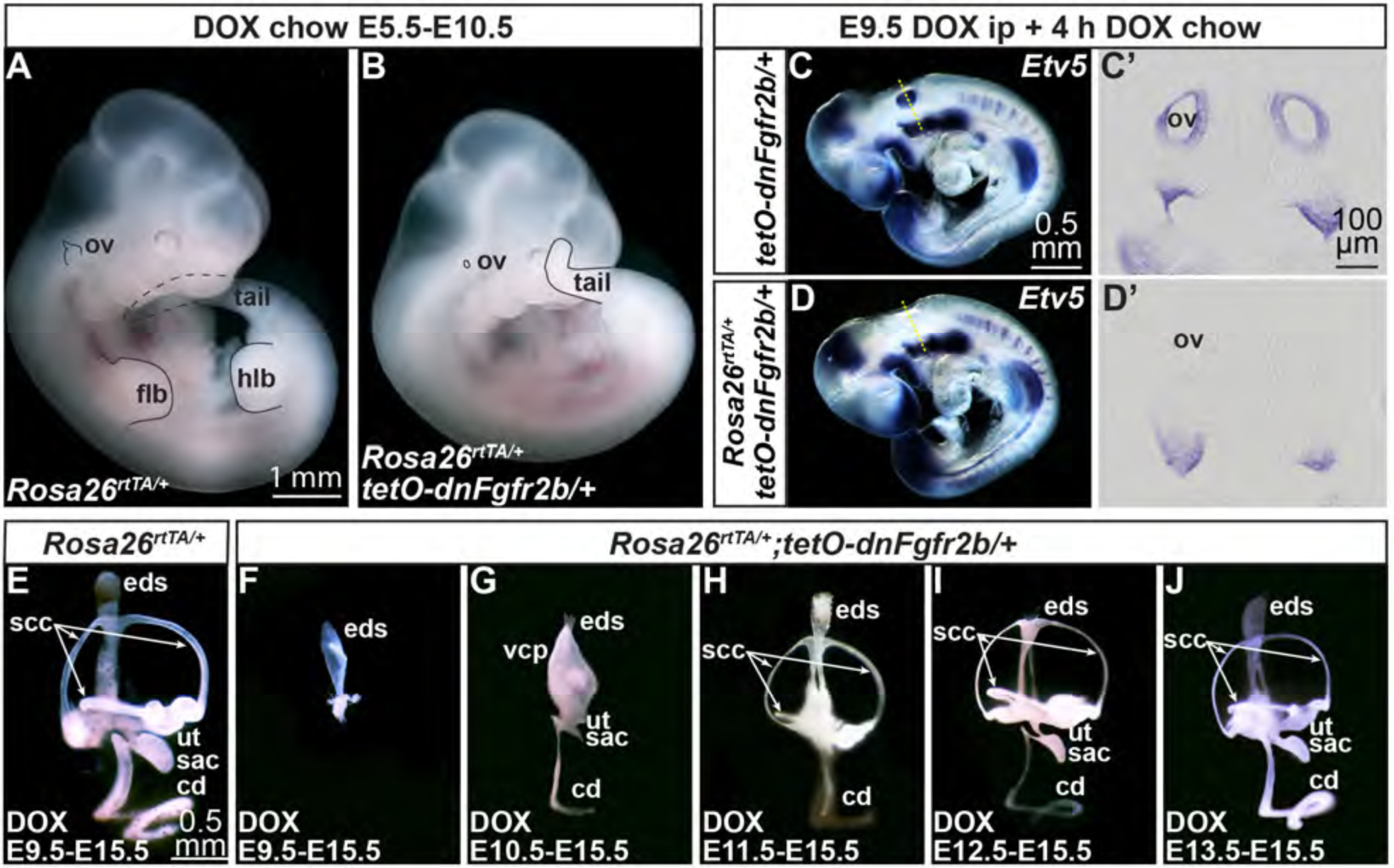
Induction of dnFGFR2b phenocopies F3KO;F10KO mutants and acts rapidly, revealing continuous requirements for FGFR2b ligands in otic morphogenesis. (A,B) E10.5 control and experimental embryos exposed to DOX from E5.5-E10.5. Affected structures outlined in black; dotted lines indicate regions behind the embryo. Scale bar in A applies to B. (C-D’) E9.5 control and experimental embryos exposed to DOX for 4 hours and hybridized with *Etv5*. Dashed lines in C and D indicate transverse section planes shown in C’,D’. Scale bars in panels C,C’ apply to D,D’. (E-J) E15.5 control and experimental paintfilled inner ears from embryos exposed to DOX for intervals indicated at the bottom of each panel. Scale bar in E also applies panels F-J. Genotypes used for each analysis are boxed. Abbreviations not used previously: eds, endolymphatic duct/sac; flb, forelimb bud; hlb, hindlimb bud; scc, semicircular canal; ut, utricle; vcp, vertical canal pouch.

### Secreted dnFGFR2b acts rapidly to inhibit signaling by FGFR2b ligands

To determine the timing of signaling inhibition we initiated dnFGFR2b expression by injecting DOX at different stages, providing DOX-chow for various intervals, and assaying for *Etv5* expression by ISH. After only 4 hours of DOX starting at E9.5, *Tg(tetO-dnFgfr2b)/+* embryos showed robust *Etv5* expression in numerous sites of FGF signaling (Fig. 4C), including throughout most of the otic vesicle (Fig. 4C’). In contrast, double heterozygotes retained many sites of *Etv5* expression, but lacked any otic *Etv5* (Fig. 4D,D’). In addition, 6 hours of DOX starting at E8.25 caused a near ablation of *Etv5* throughout double heterozygotes, including in the otic cup (Fig. S3A,A’,B,B’) and 4 hours of DOX starting at E10.25 significantly downregulated *Etv5* in the dorsolateral quadrant of the otic vesicle (Fig. S3C,C’,D,D’). Thus, inhibition of FGFR2b ligands has a rapid onset in otic tissue, consistent with studies of dnFGFR2b induction in the limb (Danopoulos et al., 2013).

### FGFR2b ligands are required continuously for otic morphogenesis

Next, we asked when FGFR2b ligands are required for otocyst morphogenesis. Starting between E8.5 and E13.5 we injected DOX into pregnant dams and provided DOX-chow continuously through E15.5 when the inner ears were paintfilled. We compared *Rosa26^rtTA/+^* (control) to *Rosa26^rtTA/+^;Tg(tetO-dnFgfr2b)/+* (experimental) samples. DOX exposure from E8.5-E15.5 inhibited inner ear development such that experimental embryos had no otic tissue to fill (n=6/6; data not shown). When DOX was started on E9.5, control ears appeared normal (Fig. 4E) and experimental samples showed two distinct phenotypes: either no detectable inner ear (n=6/10; not shown) or a structure resembling an EDS (n=4/10; Fig. 4F). Starting DOX on E10.5 blocked development of distinct SCCs from the vertical canal pouch, reduced the size of the saccule and utricle, and caused a dramatic shortening and narrowing of the CD (n=16/18; Fig. 4G). This phenotype resembled the most strongly affected F10KO mutants (Urness et al., 2015). DOX exposure from E11.5-E15.5 was compatible with SCC formation, though these appeared thinner than those of control ears (n=8/8; Fig. 4H). Development of the utricle, saccule and CD were somewhat variable, but not markedly better than in the E10.5-E15.5 samples. Inner ears exposed to DOX from E12.5-E15.5 had a much more normal morphology, but still, the SCCs and CD were narrow (n=8/8; Fig. 4I). Even experimental ears exposed to DOX from only E13.5-E15.5 showed the thin SCC defect, but the rest of the inner ear appeared grossly normal (n=8/8; Fig. 4J). These data show that FGFR2b ligands are required continuously during otic morphogenesis.

### Transient activation of dnFGFR2b reveals critical periods for FGFR2b ligands in otic morphogenesis

To determine more precise intervals for FGFR2b ligand requirements in particular events of otic morphogenesis, we treated pregnant dams with different DOX pulses and examined E15.5 inner ears by paintfilling. As expected, all ears exposed to DOX for even the longest pulse (24 h) had normal morphology (Figs. 5A,E,G,K,M,Q). In contrast, experimental ears showed exposure time-dependent abnormalities. A 4-hour DOX exposure starting at 9 AM on E8.5 (termed E8.25 for convenience) caused mild PSCC reductions in a few ears (n=3/14; Fig. 5B) and had no discernable effect on the remaining ears, whereas 6-hour or 24-hour exposures had increasingly severe consequences. Most of the 6-hour group (n=14/19) and some of the 24-hour group (n=2/16) lacked an EDS (Figs. 5C’,D). The majority of 24-hour exposures blocked most development of the otocyst, leaving a small vesicle (n=6/16; Fig. 5D’) or no ear tissue (n=8/16; data not shown). By delaying DOX administration to 9 PM on E8.5 (called E8.75) all experimental samples that had any ear tissue (n=17/32) showed at least an EDS and most (n=12/32) had a central (vestibular) segment and a linear CD (Figs. 5F,F’). A 2-hour DOX pulse starting at 9 AM on E9.5 (called E9.25) had no effect on morphogenesis (data not shown), but 4- or 6-hour exposures consistently caused only PSCC defects (Figs. 5H,I,I’). The 24-hour exposures allowed EDS outgrowth, but consistently blocked most vestibular and cochlear outgrowth (n=40/40; Figs. 5J,J’). 12-hour DOX exposures starting at 9 PM on E9.5 permitted EDS outgrowth and formation of at least a vertical canal pouch, but no SCC formation. In addition, the CD was short and narrow (n=14/16) or not present (n=2/16; Fig. 5L,L’). DOX exposures starting at 9 AM on E10.5 (termed E10.25) and extending for 4 or 6 hours consistently blocked normal PSCC formation (n=19/20; both groups considered together) and the 6-hour exposure sometimes affected the ASCC as well (n=3/10), but had virtually no effect on development of the utricle, saccule or CD (Figs. 5N,O,O’) until the exposure reached 24 hours (Figs. 5P,P’). By starting DOX at 9 PM on E10.5 (termed E10.75), the most severe defects were avoided, nevertheless, the SCCs appeared thin, the utricle and saccule were reduced and the CD was short (n=5/5; Fig. 5R). In summary, we found that the earlier DOX was started and the longer it was present, the more severe were the morphogenesis defects. Furthermore, some of the DOX pulses gave such consistent outcomes that it seemed possible to identify acute epithelial transcriptional targets of FGFR2b ligands mediating particular morphogenetic events.

**Figure 5.**
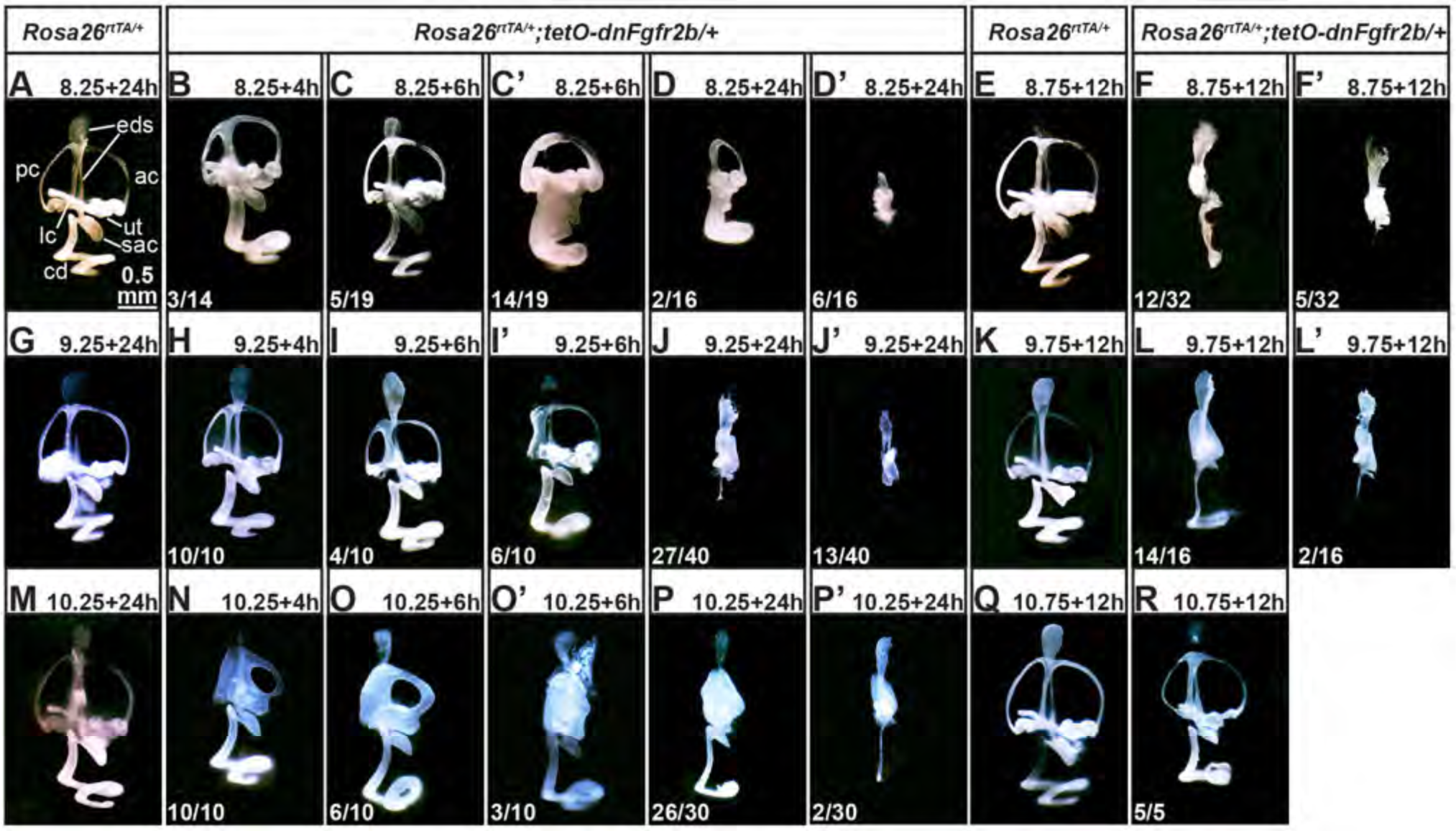
Transient activation of dnFGFR2b reveals critical periods for FGFR2b ligands in otic morphogenesis. Paintfilled ears from control and experimental embryos with genotypes boxed above. DOX was provided to pregnant dams at E8.25 (A-D’), E8.75 (E-F’), E9.25 (G-J’), E9.75 (K-L’), E10.25 (M-P’) or E10.75 (Q-R) for the hours indicated above each panel. The number of ears from each treatment showing the same phenotype is indicated at the lower left. Scale bar in A applies to all panels. Abbreviations not used previously: ac, anterior canal; lc, lateral canal; pc, posterior canal.

### RNA-Seq reveals transcriptional targets of FGFR2b ligands during early otic morphogenesis

To identify transcriptional targets of FGFR2b ligands during early otocyst morphogenesis, when they are required for both vestibular and cochlear outgrowth, we chose three DOX exposures (Fig. 6A) that gave similar morphogenesis outcomes: E9.75+12 hours (Seq1, Figs. 5L,L’), E10.25+24 hours (Seq2, Figs. 5P,P’) and E9.25+24 hours (Seq3, Figs. 5J,J’). Immediately following DOX exposure, otocysts were microdissected, cleaned of mesenchyme (Fig. 6B), and pooled into separate control and experimental groups from each female. RNA was isolated, processed for RNA-Seq and analyzed for differential expression under both unpaired (genotype only) and paired (genotype and litter) statistical models.

**Figure 6.**
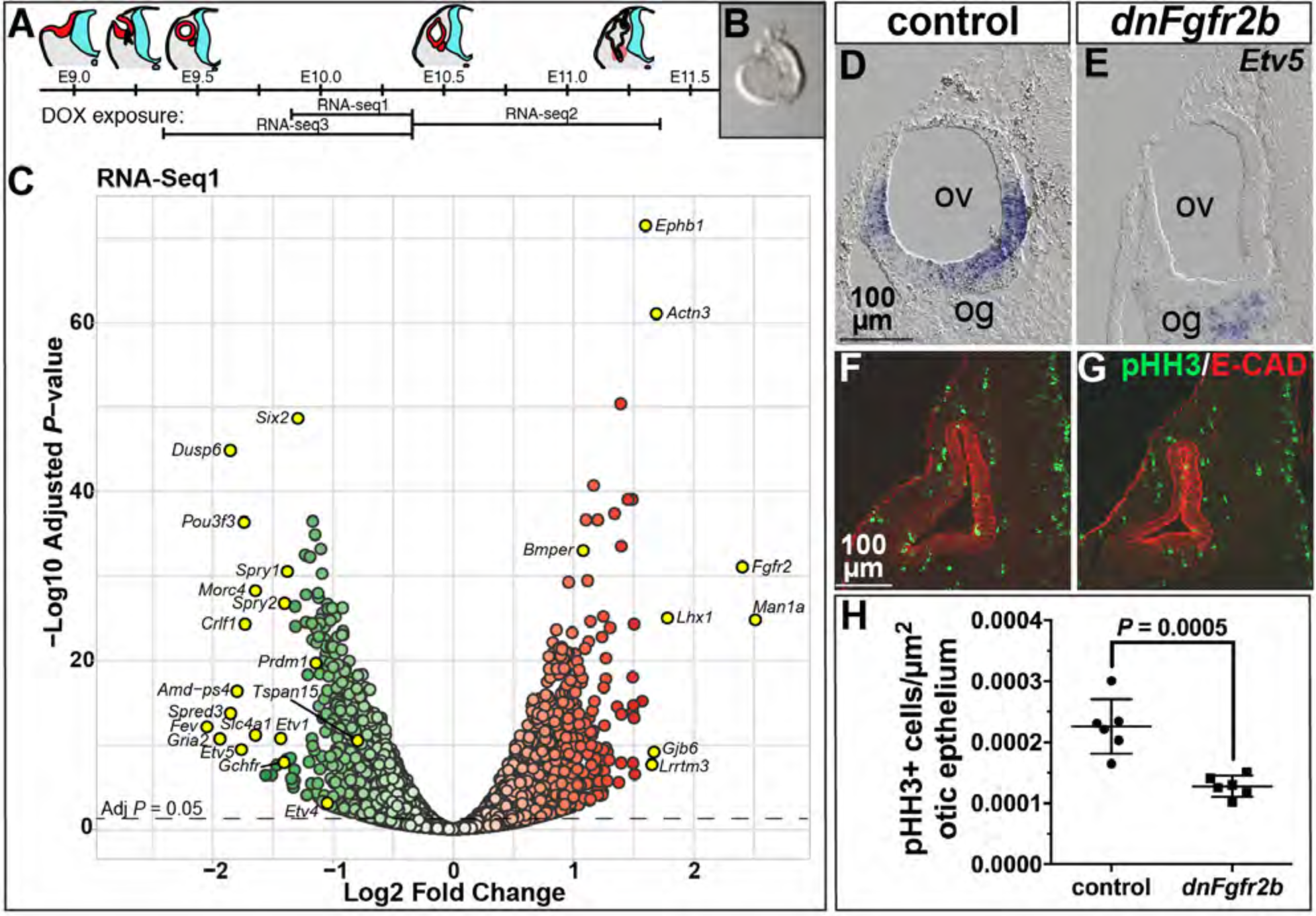
Differential RNA-Seq reveals expected and novel targets of FGFR2b ligands during early otic morphogenesis and their requirement in otic epithelial proliferation. (A) Schematic of otocyst morphogenesis correlated with the three DOX exposures used for RNA-Seq. (B) Microdissected E10.5 otocyst used for RNA isolation. (C) Volcano plot for RNA-Seq1 showing significantly downregulated (green) and upregulated (red) genes identified using a paired statistical model. Significance plotted on the y-axis and fold change on the x-axis. Gene labels highlighted in yellow indicate fold-change >1.5, common FGF target genes, and genes pursued for ISH validation. (D-E) Transverse sections of E10.25 control and *dnFgfr2b* otocysts from RNA-Seq1 embryos hybridized with a probe for *Etv5*. (F-G) Transverse sections of E10.25 control and *dnFgfr2b* otocysts from RNA-seq3 embryos immunostained for pHH3 (green) and E-Cadherin (red). (H) Quantification of pHH3-positive cells per otic epithelial area in Seq3 control and *dnFgfr2* otocysts. Scale bars in D,F apply to E, G. Abbreviations as defined previously.

Differentially expressed genes with an adjusted (adj) P<0.05 in each paired dataset were visualized with volcano plots (Fig. 6C; Fig. S4). In all 3 datasets the maximum fold-changes were relatively modest, perhaps reflecting the short periods of inhibition, but for many genes the differences were highly significant. *Fgfr2* and *Ighg1* were among the most highly differentially expressed Seq1 genes (5.3-fold and 633-fold induced, respectively, but *Ighg1* was omitted from the plot for legibility; Fig 6C). Inspection of *Fgfr2* reads showed upregulation was due to expression of the *Fgfr2b* splice isoform specifically in transgene-containing samples. The *Ighg1* reads were also transgene-specific, thus validating the efficacy of the inductions. Excluding *Fgfr2* and *Ighg1* as vector-specific, there were 968 genes >1.5-fold upregulated and 631 genes >1.5-fold downregulated (adjP<0.05) in experimental otocyst RNA. Significantly downregulated genes included well-known transcriptional targets of FGF signaling including *Etv1, Etv4, Etv5, Dusp6, Spry2* and *Spry1* (indicated in Fig. 6C and listed, Excel S1). Similar analyses of Seq2 and Seq3 also showed significant upregulation of vector-specific sequences and significant downregulation of known FGF target genes (Fig. S4, listed in Excel S1). To validate an FGF target gene significantly downregulated in all 3 datasets, we detected *Etv5* by ISH of otocyst sections. Seq1 control otocysts showed lateral and ventromedial *Etv5* expression, whereas experimental embryos did not express otocyst *Etv5* (Fig. 6D,E). Similar results were obtained with Seq3 samples (Fig. S5).

*Fgfr1* sequence reads in each dataset showed no changes in level between control and experimental samples, which were similar to control levels of *Fgfr2*. Interestingly, *Fgfr1c*, thought to be mesenchymal, was the predominant splice isoform, but *Fgfr1b* was also detected. *Fgfr3* sequences were present at levels at least 20-fold below those of *Fgfr1* or *Fgfr2* in control samples and were unchanged by dnFGFR2 induction (Excel S1). The only FGFR2b ligand genes expressed at significant levels in control or experimental otocysts were *Fgf3* and *Fgf10* (Excel S1), consistent with ISH surveys (Wright et al., 2003 and data not shown). Interestingly, *Fgf3* was slightly, but significantly upregulated in all 3 datasets. However, as dnFGFR2b inhibition acts at the level of protein, this is unlikely to impact the phenotypes.

### FGFR2b ligands are required to promote otocyst cell proliferation

To explore functional relationships between significantly differentially expressed genes in each dataset (either up or downregulated; adj*P*<0.05), we used Ingenuity Pathway Analysis. In each case, the top 5 affected pathways included cell cycle and DNA damage/repair pathways (Excel S2; -log(*P*-value)=8-13). In most cases these genes were downregulated in our datasets. In addition, we used GOrilla software to identify gene ontology terms for processes enriched in the downregulated Seq1 dataset (adj*P*<0.05), and the results were similar (top 5 shown in Excel S2; FDR q-values 1.55x10^-21^-7.84x10^-13^). To assess proliferation more directly, we quantified pHH3-positive cells per otic epithelial area in Seq3 otocyst sections and found a significant ~1.8-fold decrease in pHH3 labeling of experimental samples relative to controls (Fig. 6F-H), suggesting that one role for FGFR2b ligands during E9.25-E10.25 is to control the rate of otic epithelial proliferation.

### Signaling by FGFR2b ligands in the early otocyst represses genes that function later in otic epithelial development or hearing

Among the significantly upregulated genes in *dnFgfr2b* samples from the Seq1 paired analysis, we noticed a gene that, when mutated, causes human hearing loss *(Gjb6;* Fig. 6C). To determine whether other such genes, or those responsible for mouse hearing loss and/or cochlear development, were enriched in either the up- or downregulated genesets, we conducted a gene set enrichment analysis (Subramanian et al., 2005) on all 16,232 genes detected in the Seq1 paired analysis using two partially overlapping gene lists: 95 human hereditary hearing loss genes identified by Nishio et al. (2015; rank listed in Excel S3) and 258 mouse genes involved in inner ear development or function collated by Ohlemiller et al. (2016; rank listed in Excel S4). Both genesets were highly enriched in the upregulated Seq1 dataset (normalized enrichment score=2.09 for the human genes and 2.06 for the mouse genes; both nominal *P*-values, <1x10^-3^), but not in the downregulated set. Similar results were obtained with the Seq2 dataset (data not shown). Together, these analyses suggest that one role of FGFR2b ligands at this early stage of morphogenesis is to prevent premature expression of epithelial genes that have later roles in development or function of the inner ear.

### Validation of new genes regulated by FGFR2b ligands during the early stages of otocyst morphogenesis

To validate new FGFR2b target genes by ISH we focused first on downregulated genes and assayed selected genes based on overlap in multiple RNA-Seq datasets (Excel S5), relatively high degree of differential expression, and normalized read count above that of *Fgf3* (Excel S1), which is relatively difficult to detect by ISH, and novelty with respect to inner ear development and/or FGF/MAPK signaling. Seven genes validated as downregulated in Seq1 otocysts are illustrated in Figure 7. *Spred3* was expressed in E10.25 control otocysts in a ventromedial domain (Fig. 7A), and was greatly reduced in experimental otocysts (Fig. 7A’). *Six2* was detected in the ventral-most region of control otocyts as well as laterally (Fig. 7B), but was absent from dnFGFR2b otocysts (Fig. 7B’). *Prdm1(Blimp1)* was expressed similarly to *Six2* in control otocysts (Fig. 7C) and was strongly downregulated in the corresponding experimental otocysts (Fig. 7C’). *Crlf1* was also expressed similarly to *Six2* (Fig. 7D) in controls, but ventral expression was absent and lateral expression was downregulated in dnFGFR2b otocysts (Fig. 7D’). *Tspan15* was expressed in a broad ventrolateral domain in controls (Fig. 7E) and was largely absent from dnFGFR2b otocysts (Fig. 7E’). *Pou3f3* was expressed in the ventral-most region of controls (Fig. 7F) and was absent from dnFGFR2b otocysts (Fig. 7F’). Finally, *Gchfr* was expressed similarly to *Pou3f3* (Fig. 7G) and was absent from dnFGFR2b otocysts (Fig. 7G’). We observed similar downregulation of *Spred3, Prdm1, Crlf1, Tspan15*, and *Gchfr* expression in dnFGFR2b otocysts subjected to the Seq2 and Seq3 DOX exposures (Figs. S6,S7). *Six2* and *Pou3f3* downregulation was confirmed by ISH in Seq3 otocysts (Fig. S7), but not tested in Seq2 otocysts, as these genes were not significantly affected in the Seq2 dataset.

**Figure 7.**
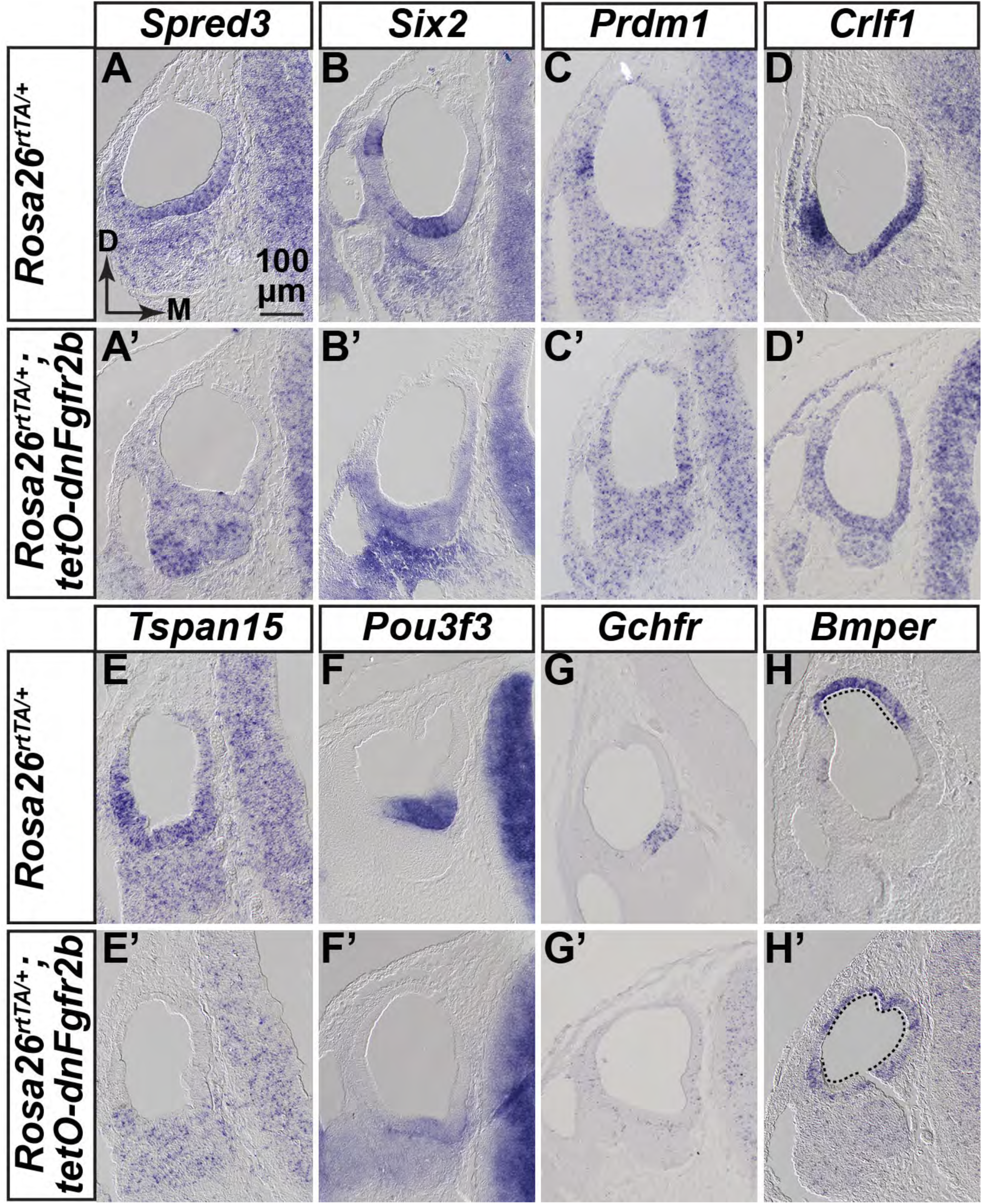
Validation of new genes regulated by FGFR2b ligands during early otocyst morphogenesis. ISH of transverse sections of E10.25 RNA-Seq1 control and *dnFgfr2b* otocysts (n=3 each). Probes are indicated above each column and genotypes are indicated to the left of each row. Scale and orientation for all panels indicated in A.

We also tested several genes common to the upregulated lists, but most were widely expressed in controls and any changes in expression levels were not revealed by ISH (data not shown). However, *Bmper* transcripts were confined to the dorsomedial region of control otocysts (Fig. 7H), but the expression domain expanded to encompass most of the otic epithelium in experimental otocysts (Fig. 7H’). Similar results obtained with Seq2 and Seq3 otocysts (Figs. S6,S7). Therefore, these datasets are a rich source of novel FGFR2b signaling targets in the otocyst.

## DISCUSSION

*Fgfr2b* and two genes encoding FGFR2b ligands, *Fgf3* and *Fgf10*, are required individually for otic morphogenesis (Hatch et al., 2007; Mansour et al., 1993; Pauley et al., 2003; Pirvola et al., 2000; Urness et al., 2015). We found that *Fgf3* and *Fgf10* are expressed continuously during otic morphogenesis, however, their requirement for otic placode induction (Alvarez et al., 2003; Urness et al., 2010; Wright and Mansour, 2003a) obscured potential combinatorial roles during otocyst morphogenesis. Here, we blocked FGF3 and FGF10 signaling after otic placode induction by using conditional gene inactivation and temporally controlled inhibition via a drug-inducible ligand trap. Analysis of conditional mutants revealed that both genes are required in the otic lineage after E8.5 for both vestibular and cochlear development, but that the role of *Fgf3* is only revealed in the absence of *Fgf10*, and that these signal-encoding genes control expression of genes that function in otic epithelial morphogenesis. Furthermore, we found that FGF3 and FGF10 signal redundantly to maintain otic neuroblasts. Temporally regulated inhibition showed that FGFR2b signaling acts continuously throughout otic morphogenesis and that pulses of inhibition can be used to identify the timing of particular morphogenetic events. We used brief windows of inhibition at early stages of morphogenesis to conduct the first genomewide identification of proximal targets of FGFR2b signaling in the otocyst, and validated several genes that are novel candidates for involvement in the initial stages of otocyst morphogenesis.

### *Fgf3* and *Fgf10* are expressed and function continuously to control otocyst formation and morphogenesis

*Fgf3* and *Fgf10* are expressed continuously throughout otocyst development and cochlear morphogenesis. These are likely the only relevant FGFR2b (or FGFR1b) ligand-encoding genes for early otic morphogenesis as, based on RNA-Seq data, the others are either not expressed *(Fgf22)* or are detected at negligible levels *(Fgf1* and *Fgf7)*. The use of stage-matched tissue throughout morphogenesis helped to define the relative temporal and spatial dynamics of the expression patterns and aided in interpreting functional perturbations. We found that *Fgf10* has an earlier and broader distribution than *Fgf3* in the otic epithelium,and both transcripts are present in the otic ganglion, but *Fgf3* is seen only transiently, whereas once *Fgf10* starts expressing, it is present continuously at high levels in the cochlear ganglion. Although *Fgf10* is expressed in mesenchyme underlying the preplacodal ectoderm, neither gene appears in periotic mesenchyme during otic cup formation or later (Schimmang, 2007; Urness et al., 2015; Wright and Mansour, 2003b). ISH data for *Fgfr2b* and *Fgfr1b*, which encode the receptors for FGF3 and FGF10, are limited because of the small size of probes that distinguish them from “c” isoforms, but extant data are consistent with the idea that they are also primarily epithelial (Orr-Urtreger et al., 1993; Pirvola et al., 2000; Wright et al., 2003; Wright and Mansour, 2003a) and, indeed, the “b” isoforms are evident in our otocyst RNA-Seq datasets. Therefore, taken together, the ligand and receptor expression data are consistent with our findings of continuous and combinatorial roles for *Fgf3* and *Fgf10* in otocyst epithelial morphogenesis and otic ganglion development. They also raise the interesting possibility that signaling involves epithelial and/or ganglion ligands activating epithelial, rather than mesenchymal or ganglionic receptors, except perhaps as neuroblasts begin delamination from the epithelium. This contrasts with epithelial ligands FGF9 and FGF20, which signal initially to “c” type receptors in the mesenchyme, inducing signals that control epithelial proliferation (Huh et al., 2015).

### *Fgf3* and *Fgf10* are both required for vestibular and cochlear morphogenesis and for maintenance of otic neuroblasts

Conditional mutant analyses showed that both genes are required after otic placode induction, but deletion of *Fgf3* alone from the *Pax2-Cre* lineage was inconsequential. This contrasts with the variably penetrant otic dysmorphology of F3KO mutants, the most severe of which initiate with alterations of dorsal otocyst patterning, loss of the EDS, and subsequent cystic development of the epithelium, ultimately resulting in hearing loss and circling behavior (Hatch et al., 2007; Mansour et al., 1993). The normal phenotype of F3cKO ears points to a critical role for *Fgf3* expression in the hindbrain. Indeed, we found that F10KO embryos in which only hindbrain sources of *Fgf3* were deleted (using *Sox1^Cre^)* had very small otocysts. In contrast, F10cKO ears had abnormalities very similar to those of F10KO ears. This demonstrates that the unique functions of *Fgf10* in otic morphogenesis arise from its expression in the placodal lineage rather than earlier in the mesenchyme. Analysis of conditional mutants that separate epithelial from ganglion sites of *Fgf10* expression will be needed to further dissect the spatial requirements for *Fgf10* function.

Although *Pax2-Cre* is active in the placode at E8.5, we found no obvious effects at E9.5 on otocyst morphology in F3cKO;F10cKO embryos, and only two tested genes, *Etv5* and *Tbx1*, were lost or altered in these otocysts. The first major losses in expression of multiple genes required for morphogenesis occurred at E10.5. This shows that both *Fgf3* and *Fgf10* are required in the placode lineage for normal otocyst morphogenesis and suggests that overlapping expression of *Fgf3* and *Fgf10* starting at E9.5 may be critical for both cochlear and vestibular outgrowth and morphogenesis. However, the phenotypes of triple allelic conditional mutants point to functional differences between *Fgf3* and *Fgf10*. Both the cochlear and vestibular morphology of F3cKO/F10cHet ears were less severely affected than in F3cHet/F10cKO ears. This may reflect the relatively larger domain and higher level of epithelial *Fgf10* than of *Fgf3*, and may be presaged by the differential effects on *Etv5* expression in the two types of E9.5 otocysts. The loss of dorsolateral *Etv5* when *Fgf10* is the only remaining allele, and of ventromedial *Etv5* when *Fgf3* is the only remaining allele, suggest that *Fgf3* is particularly important dorsally and *Fgf10* ventrally, at least initially. The loss of dorsolateral *Tbx1* in the two most severely affected genotypes likely reflects effects of FGF3/FGF10 signaling on development of the vertical canal pouch, the derivatives of which (PSCC and ASCC) are strongly affected in these and in *Tbx1* mutants (Freyer et al., 2013; Macchiarulo and Morrow, 2017). Whether this is a direct or indirect effect on *Tbx1* expression is not yet clear, but it is interesting to note that *Tbx1* is slightly, but significantly downregulated in the Seq2 and Seq3 datasets (Excel S1).

The presence of an EDS in both triple allelic conditional mutants and normal *Gbx2* expression in these mutants and in F3cKO/F10cKO mutants contrasts with findings from F3KO mutants (Hatch et al., 2007), which usually lack an EDS and lose *Gbx2* expression by E10.5. This is consistent with the idea that hindbrain, rather than epithelial *Fgf3*, induces the EDS. It is possible that the further shortening of F3cHet/F10cKO CDs results from reduced FGF3-stimulated proliferation rather than alterations in molecular patterning. This is supported by preliminary analyses of E18.5 ears that did not reveal any exacerbation of changes to CD marker genes analyzed previously in the F10KO mutant (data not shown, see Urness et al., 2015). However, the timing of such proliferative effects in F3cKO;F10cKO mutants must be later than E10.5, when differences in pHH3 labeling between control and F3cKO/F10cKO otocysts were not significant.

We suggested previously that *Fgf3* plays a role in otic ganglion development, as the F3KO ganglion, like that of *Fgfr2b* null mutants (Pirvola et al., 2000), is smaller than normal (Mansour et al., 1993). In contrast, F10KO early otic ganglia and later cochlear ganglia appear normal (Urness et al., 2015) despite defects of vestibular innervation consequent to midgestation loss of vestibular sensory epithelia (Pauley et al., 2003). In contrast to zebrafish (Vemaraju et al., 2012), our present data from F3cKO;F10cKO mutants suggest that *Fgf3* and *Fgf10* are required together for maintenance of *Neurog1* and *Neurod1* expression, and development of an otic ganglion, rather than specification of otic neuroblasts. Our data do not address whether this requirement involves ligand expression in the epithelium or ganglion or both. However, by restricting dnFGFR2b expression to the placodal lineage by using *Pax2-Cre* in combination with the unrecombined *Rosa26^lslrtTA^* allele, it may be possible to avoid disrupting otic induction and determine whether FGFR2b ligands play any role in mouse otic neuroblast specification. In addition, this paradigm of tissue restricted and timed induction of dnFGFR2b could also be used to identify candidate genes responsible for neuroblast maintenance. Determining the role of the transient burst of *Fgf3* in delaminating neuroblasts will be more challenging.

### Temporally controlled inhibition of FGFR2b signaling during otocyst morphogenesis reveals requirements at multiple stages

Observations of embryos expressing dnFGFR2b between E5.5 and E10.5 showed that this inhibition strategy effectively phenocopies F3KO;F10KO mutants without causing abnormalities characteristic of disrupting FGFs that signal through “c”-type receptors. Thus, dnFGFR2b has the expected specificity and the only relevant FGFR2b ligands for otocyst morphogenesis are likely to be FGF3 and FGF10. Paintfilling of embryonic ears subjected to ubiquitous and chronic dnFGFR2b expression starting on different days of development revealed that FGFR2b ligands are required continuously for otic development at least through E13.5. Pulses of dnFGFR2b caused highly specific and penetrant otic malformations, supporting the idea that unlike irreversible CRE-mediated deletion of coding exons, the signaling inhibition effected by dnFGFR2b is reversible.

Together, the chronic and pulsed inhibition paradigms suggest a distinct progression of roles for FGFR2b ligands: first in inducing the placode, then in stimulating EDS, vestibular pouch and CD outgrowth, and finally in sculpting the SCCs, outgrowth of utricle and saccule, and specification of CD nonsensory tissue. As otic induction is complete by ~E8.5, it was surprising that starting chronic dnFGFR2b expression on E9.5 blocked ear development in 6/10 cases. Since dnFGFR2b embryos from Seq3 inductions (E9.25-E10.25) always had otocysts, and these had significantly reduced proliferation, we suggest that the loss of otic tissue in embryos induced chronically after E8.5 reflects degeneration of mitotically blocked cells rather than failure of otic induction. Some phenotypes revealed the timing of roles for FGFR2b ligands. For example, the 6–24 hours starting at E8.25 is particularly important for EDS formation, potentially reflecting FGF3 inhibition (Mansour et al., 1993; Hatch et al., 2007) and the 6-hours starting at E9.25 is important for PSCC formation, potentially reflecting FGF10 inhibition (Pauley et al., 2003; Urness et al., 2015). Other vestibular phenotypes are particularly interesting as they reveal three potential and previously unsuspected functions for FGFR2b signaling. Chronic dnFGFR2b induction starting at E10.5, or 4–24 hour pulses starting at E10.25 blocked fusion of vestibular pouches, suggesting requirements for FGFR2b ligands in fusion plate formation and chronic induction starting between E11.5-E13.5 or 12-hour pulses starting E9.75-E10.75 caused very thin semicircular canals, suggesting roles for FGFR2b ligands in limiting resorption of fusion plates. Finally, several conditions reduced utricle and saccule development. Thus, it will be interesting to explore regulatory relationships between FGFR2b signaling and genes already known to regulate vestibular mophogenesis (Alsina and Whitfield, 2017), as well as to induce dnFGFR2b in particular temporal/spatial windows and pursue unbiased identification of effector genes involved in the development of particular structures of interest.

### FGFR2b ligands promote otic epithelial proliferation and prevent premature expression of genes required for hearing

Our RNA-Seq datasets revealed significant downregulation of genes involved in the cell cycle and DNA repair, and indeed, immunostaining of Seq3 samples showed that mitotic cell numbers were significantly reduced in E10.25 dnFGFR2b-containing otocysts. This result differed from that obtained with E10.5 F3cKO/F10cKO otocysts, which did not show a mitotic defect. Given that hindbrain *Fgf3* is unaffected in *Pax2-Cre*;F3cKO/F10cKO mutants, and is extinguished by E10.5, it is likely that otocyst proliferation defects in these mutants would be detected at later stages of morphogenesis.

The RNA-Seq datasets also revealed significantly upregulated genes. We found that in Seq1 and Seq2, these genes are highly enriched for human hereditary hearing loss genes and mouse genes that are expressed and/or function later in the inner ear. These include *Pax2*, which was expanded in E10.5 F3cKO/F10cKO otocysts. This suggests that at early stages, FGFR2b signaling normally represses many genes important for later development and function of the cochlea, or alternatively, that the proliferative block imposed by dnFGFR2b expression promotes early differentiation of the epithelium. The latter possibility however, does not apply to the earliest genes required for sensory cell differentiation *(Atoh1, Pou4f3* and *Gfi1)*, which were detected at very low levels and were unaffected in any of the RNA-Seq datasets. We suggest that the upregulated genesets are worth mining for new candidates for hearing loss genes, of which many remain to be identified (Bowl and Brown, 2018).

Although we show that FGFR2b ligands are required to activate a *Bmp (Bmp4)* and repress a BMP regulatory gene *(Bmper)*, we found no evidence for FGFR2b ligand regulation of key downstream components of the BMP or SHH pathways, suggesting that while FGFR2b ligands may regulate individual components of these pathways, at least at the stages investigated, they are not exclusively upstream of these key programs directing dorsal and ventral otic morphogenesis, respectively.

### Novel targets of FGFR2b signaling in early otocyst development

We validated by ISH seven novel genes downregulated and one gene upregulated by FGFR2b ligands in the RNA-seq datasets. Some genes may be regulated directly by the intracellular signaling pathway activated by FGFR2b, as one of our analysis points was only 12 hours after induction. As it was not possible to study more than a few differentially expressed genes, it is difficult to speculate about their combinatorial functions in inner ear development. Nevertheless, it is interesting to note that the downregulated genes in our most robust dataset (Seq1) are enriched for transcription factor-coding genes (Excel S2), including those validated here, *Six2, Prdm1* and *Pou3f3*. The first two have otocyst expression patterns similar to *Spred3, Crlf1* and *Tspan15*, whereas *Pou3f3* expression appears to overlap with *Gchfr*. This suggests that further mining of the existing differential expression data and generation of additional targets by employing different windows of FGFR2b inhibition, combined with promoter analysis and genomewide studies of otocyst chromatin modification could suggest important new gene regulatory networks acting to shape the epithelium.

The only upregulated gene validated by ISH was *Bmper*, which encodes BMP-binding endothelial regulator, an ortholog of *Drosophila* crossveinless-2 (cv-2) (Coffinier et al., 2002). cv-2 modulates BMP signaling biphasically in the fly wing with the direction of action dependent upon the concentration of Cv-2 and the concentration and types of local BMP ligands (Conley et al., 2000; Serpe et al., 2008). Similarly, in mouse, *Bmper* modulates the availability of BMPs, enhancing signaling when ligands are low and limiting signaling when ligands are high (Dyer et al., 2014; Kelley et al., 2009). Thus, *Bmper* null mutants have some phenotypes suggestive of a classic BMP signaling antagonist (Moser et al., 2003) and others suggestive of a BMP signaling agonist (Ikeya et al., 2006). Multiple *Bmps* and their receptors are expressed in and required for otocyst morphogenesis (Chang et al., 2008; Hwang et al., 2010; Ohyama et al., 2010) and misexpression of BMP ventral to the otic placode blocks outgrowth of the chick cochlea (Ohta et al., 2016). Thus, it will be interesting to determine whether the otic phenotype of a *Bmper* null mutant reflects a loss or gain of BMP signaling, and whether this differs in different regions of the developing otocyst. Our results also showed that at early stages of otocyst morphogenesis, *Fgf3* and *Fgf10* are required for expression of *Bmp4*, which is itself required for both vestibular and cochlear development (Chang et al., 2008). Determining whether FGFR2b ligand-dependent upregulation of *Bmper* functions in this context to further antagonize BMP signaling, or alternatively, to mitigate the reduction in *Bmp4* by increasing signaling by other BMP ligands will require additional studies of otic *Bmp* expression and manipulation of *Bmper* allele levels in combination with dnFGFR2b induction at different stages.

The identification of several FGFR2b target genes not implicated previously in ear development or hearing loss syndromes provides a tantalizing glimpse into a new set of potential otocyst morphogenetic factors. Given the novelty of these targets, it is tempting to speculate that previously unappreciated regulatory pathways may be at play during otic morphogenesis, as has been postulated for otic placode induction (Anwar et al., 2017). Functional studies will be required to address the roles of each of these new genes.

## MATERIALS AND METHODS

### Mouse models and genotyping

Mice were maintained and euthanized in accordance with protocols approved by the University of Utah Institutional Animal Care and Use Committee. All *Fgf* mutant alleles were kept on a mixed genetic background comprised of C57Bl/6 and various 129 substrains. CD-1 outbred mice (Charles River Laboratory) were used to generate embryos for studies of normal expression patterns and for generating embryos for induction of dnFGFR2b. Noon of the day a mating plug was observed was considered E0.5.

Generation and PCR genotyping of the *Fgf3* and *Fgf10* null alleles (*Fgf3*^-^, formally designated *Fgf3^tm1.1Sms^ =* MGI:3767558 and *Fgf10^-^*, formally designated *Fgf10^tm1.1Sms^* = MGI:3526181) and *Fgf3* and *Fgf10* and conditional alleles (*Fgf3^c^*; *Fgf3^tm1.2Sms^* = MGI:4456396] and *Fgf10^c^*; *Fgf10^tm1.2Sms^* = MGI:4456398) were described previously (Hatch et al., 2007; Urness et al., 2010). Tg(Pax2-Cre) mice (Tg(Pax2-cre)1Akg = MGI:3046196) were obtained from Dr. Andrew Groves (Ohyama and Groves, 2004). Tg(Pax2-Cre) was detected by PCR using primers specific to the transgene (5’ GGGGATCCCGACTACAAGG 3’; 5’ TAGTGAGTCGTATTAATTTCGATAAGC 3’). The *Sox1^Cre^* allele (Takashima et al., 2007) was transferred from Dr. Mario Capecchi with permission from Dr. Shin-Ichi Nishikawa (RIKEN) and genotyped using generic *Cre* primers. *Rosa26^lslLacZ^* reporter mice *(Gt(ROSA)26Sor^tm1Sor^* = MGI:1861932) (Soriano, 1999) were maintained as homogyzotes.

Single conditional mutants were generated by crossing *Fgf3^c/c^* females to *Fgf3^-/+^;Tg(Pax2-Cre)/+* males or *Fgf10^c/c^* females to *Fgf10^-/+^;Tg(Pax2-Cre)/+* males. Combinations of *Fgf3* and *Fgf10* conditional mutants were obtained by crossing *Fgf3^c/c^;Fgf10^c/c^* females to *Fgf3^-/+^;Fgf10^-/+^;Tg(Pax2-Cre)/+* or *Fgf3^-/+^;Fgf10^-/+^; Sox1^Cre/+^* males. CRE activity was confirmed by mating males to *Rosa26^LacZR/LacZR^* females harvesting embryos at the indicated stages and staining with X-gal as described (Yang and Mansour, 1999).

The germline recombined *Rosa26^rtTA^* allele (derived from Gt(ROSA)26Sor^tm1(rtTA,EGFP)Nagy^; MGI:3583817) (Belteki et al., 2005; Parsa et al., 2008) and *Tg(tetO-dnFgfr2b)* (Tg(tetO-Fgfr2b/Igh1.3Jaw; MGI 5582625) (Hokuto et al., 2003) alleles were transferred from the laboratory of Dr. Saverio Bellusci with permission from Dr. Jeffery Whitsett (Cincinnati Children’s Medical Center). Genotyping primers to detect *Tg(tetO-dnFgfr2b)* were 5’ CAGGCCAACCAGTCTGCCTGGC 3’ and 5’ CGTCTGAGCTGTGTGCACCTCC 3’. *ROSA26^rtTA^* genotyping primers were ROSA5 (5’ GAGTTCTCTGCTGCCTCCTG 3’) and ROSA3 (5’ CGAGGCGGATCACAAGCAATA 3’), which generate a wild type band of 322 bp and ROSA5 and RTTA3 (5’ AAGACCGCGAAGAGTTTGTC 3’), which generate a 215 bp rtTA-specific product. Double heterozygotes were obtained initially by crossing single heterozygotes. For most of the studies described here, we crossed wild type CD-1 females to *Rosa26^rtTA/rtTA^;Tg(tetO-dnFgfr2b)/+* males, generating 50% each of control and experimental genotypes.

### RNA in situ hybridization

Embryos were harvested and fixed in 4% PFA and stored in methanol at -20°C. RNA ISH to whole mount embryos or paraffin-embedded sections were performed as described (Urness et al., 2008; Urness et al., 2010). Probes for *Sox9, Fgf3, Fgf10, Bmp4, Etv5, Gbx2, Sox2, Foxg1, Pax2, NeuroD1, Ngn1 and Crlf1* were generated by transcription of cDNA-containing plasmids. Template plasmids and acknowledgements are shown in Table S1. All other RNA probes were generated by transcription of a PCR-amplified, gene-specific 3’ UTR fragment containing a T7 promoter. The primer sequences are shown in Table S2. Whole embryos were photographed using a stereomicroscope (Zeiss Discovery.V12) fitted with a digital camera (QImaging Micropublisher 5.0). Hybridized tissue sections were photographed under DIC illumination (Zeiss Axioskop) using a digital camera (Zeiss Axiovision or Lumenera Infinity3).

### Immunostaining of frozen tissue sections for quantification of mitotic cells in the otocyst

Embryos were fixed in 4% paraformaldehyde solution and cryosectioned in the transverse plane for immunostaining as described (Urness et al., 2015). Rabbit anti-phosphohistone H3 (Millipore 06-570) was applied at a dilution of 1:400 and mouse monoclonal anti-E-cadherin (BD Biosciences 610181) was diluted 1:60. Secondary antibodies were from Invitrogen and diluted 1:400 into PBST/5% normal serum (Alexa Fluor^®^ 488 goat anti-rabbit (A11034) and Alexa Fluor^®^ 594 goat anti-mouse (A11032)). DAPI was included in the mounting medium (Vectashield, Vector Labs). Fluorescent signals were observed under epi-illumination on a Zeiss Axioskop and captured using an Infinity3 camera (Lumenera) driven by InfinityAnalyze software. Channels were overlaid using Photoshop CS5. All pHH3-positive cells in the otocysts (defined by E-Cadherin staining) were counted from 6 μm *(Pax2-Cre* cross) or 8 μm *(dnFgfr2b* cross) sections extending from anterior to posterior. N=8 control (either *Fgf3^-/c^;Fgf10^-/c^* or *Fgf3^c/+^;Fgf10^c/+^;Pax2-Cre/+)* and n=6 experimental *(Fgf3^-/c^;Fgf10^-/c^;Pax2-Cre/+)* samples were counted for the *Fgf3/Fgf10/Pax2-Cre* conditional cross. N=3 control *(Rosa26^rtTA/^+)* and n=3 experimental *(Rosa26^rtTA/+^;tetO-dnFgfR2b/+)* samples for the dnFGFR2b cross. pHH3-positive cells per ear were normalized to the cross-sectional area counted. Statistical significance was determined using an unpaired Student’s t-test (Prism software 7.0).

### Paint filling of embryonic inner ears

Filling of embryonic inner ears with latex paint and photography was as described previously (Urness et al., 2015).

### Induction of dnFGFR2b expression

Initial inductions of dnFGFR2b designed to phenocopy *Fgf3/Fgf10* double mutants were achieved by feeding pregnant females DOX chow (200 mg/kg, Custom Animal Diets, LLC) ad libitum for the indicated time periods (E5.5-E10.5, E5.5-E11.5 or E6.5-E11.5). All subsequent inductions to generate samples for paintfilling, RNA-seq, ISH or immunostaining were initiated by a single intraperitoneal injection of the pregnant dam with 0.1 ml/10 g body weight of 0.15 mg/ml (1.5mg/kg body weight) doxycycline hyclate (Sigma-Aldrich) prepared in PBS followed by provision of DOX chow ad libitum for the indicated time periods. We avoided using female *Rosa26^rtTA^* parents, as these seemed to require larger and variable amounts of DOX to see phenotypes than when rtTA was contributed by the male parent, presumably because the widespread, ubiquitous expression of rtTA in females served to sequester DOX. We did not measure the time needed to reactivate signaling after DOX withdrawal, but based on studies of the limb (Danopoulos et al., 2013), we expect that signaling resumes after 12-24 hours.

### Otic vesicle preparation and RNA isolation

Embyos from timed matings of CD-1 females and *Rosa26^rtT/ArtTA^;Tg(tetO-(s)dnFgfr2b)/+* males, with DOX exposures as specified, were dissected and the yolk sacs saved for genotyping. The otic vesicles, including surrounding mesenchyme, were crudely dissected from the head. Isolation of the vesicles free of mesenchyme was accomplished similarly to methods previously described (Urness et al., 2010) with the following modifications. Otocysts with adherent mesenchyme were incubated in 50 μl ice-cold PT solution (25 mg/ml pancreatin (Sigma), 5 mg/ml trypsin (Sigma), and 5 mg/ml polyvinylpyrrolidone MW360 (Sigma) in Tyrode’s solution) for ~7 min. (E10.25), or ~8 min. (E11.25) to promote separation of the mesenchyme. Otocysts were aspirated to Hepes-DMEM-10% FBS, where the digested mesenchyme could be gently teased from the underlying epithelium using fine forceps or tungsten needles, and by “rolling” the vesicle over the bottom of the dish to detach the mesenchyme as it adhered to the plastic. The two otocysts from each embryo were aspirated into 100 μl RNALater (Ambion) and stored at -20°C prior to genotyping. For each of four pregnant females per DOX induction regime, all otocysts of the same genotype were combined into paired control *(Rosa26^rtTA/+^)* and experimental *(Rosa26^rtTA/+^;Tg-(tetO-(s)dnFgfr2b)/+)* pools (n = 6-12 otocysts/pool).

Total RNA from each control and experimental otocyst pool was prepared using a Micro RNAeasy kit (Qiagen #74004) and analyzed for quantity and quality on a BioAnalyzer RNA TapeStation. All 24 samples (2 genotypes x 4 females x 3 DOX exposures) exceeded a RIN quality control number of 8.

### RNA-Seq and bioinformatics

RNA library preparation, sequencing and analyses were conducted by the University of Utah/Huntsman Cancer Institute High-Throughput Genomics and Bioinformatic Analysis Shared Resource. Each RNA library was prepared using a TruSeq Stranded mRNA Sample Prep kit (Illumina) with oligo(dT) selection. 50-cycle single read sequencing of each library was conducted on an Illumina Hi-Seq 2500. Sequencing reads were aligned to mm10 + splice junctions (Ensembl build 74) using Novoalign (v2.08.03). Spliced alignments were converted back to genomic space, sorted and indexed using USeq (v8.8.8) SamTranscriptomeParser. Normalized coverage tracks (coverage per million mapped reads) were generated using USeq Sam2USeq and USeq2UCSCExe. Read counts for each gene were generated using USeq DefinedRegionDifferentialSeq (Nix et al., 2008) and differential expression analysis was performed using DESeq2 (Love et al., 2014).

To inspect *Fgfr* splicing we merged each set of control and *dnFgfr2b* alignments to separate .bam files, uploaded them to IGV 2.4.10 (Robinson et al., 2011; Thorvaldsdottir et al., 2013) and generated Sashimi plots. To identify significantly regulated pathways (P<0.05, Fisher’s Exact Test), all differentially expressed genes were loaded into Ingenuity Pathway Analysis (QIAGEN Inc., https://www.qiagenbioinformatics.com/products/ingenuitypathway-analysis). For GSEA analysis, two custom gene sets based on human hearing loss genes from Nishio et al. (2015) and mouse inner ear genes from Ohlemiller et al. (2016) were loaded into the Broad Institute GSEA website (Subramanian et al., 2005) and compared to ranked lists of otocyst genes sorted by fold change from DESeq2.

## ACKNOLWLEGMENTS

We thank Katia Hatch for auditory testing of F3cKO mice, Leslie Slota for unpublished work on marker gene expression in F3cHet/F10cKO cochleae, Saverio Bellusci and Denise Al-Alam for transferring *Rosa26^rtTA^* and *Tg(tetO-dnFgfr2b)* mice, Mario Capecchi for transferring *Sox1^Cre^* mice, Tim Mosbrugger and Chris Stubben for bioinformatics support, Shannon Odelberg for help with Prism and Gary Schoenwolf for experimental and editorial advice.

## DATA AVAILABILITY

The three RNA-seq datasets are deposited in GEO under accession #GSE116404.

## Supplemental Table Legends

**Table S1:**
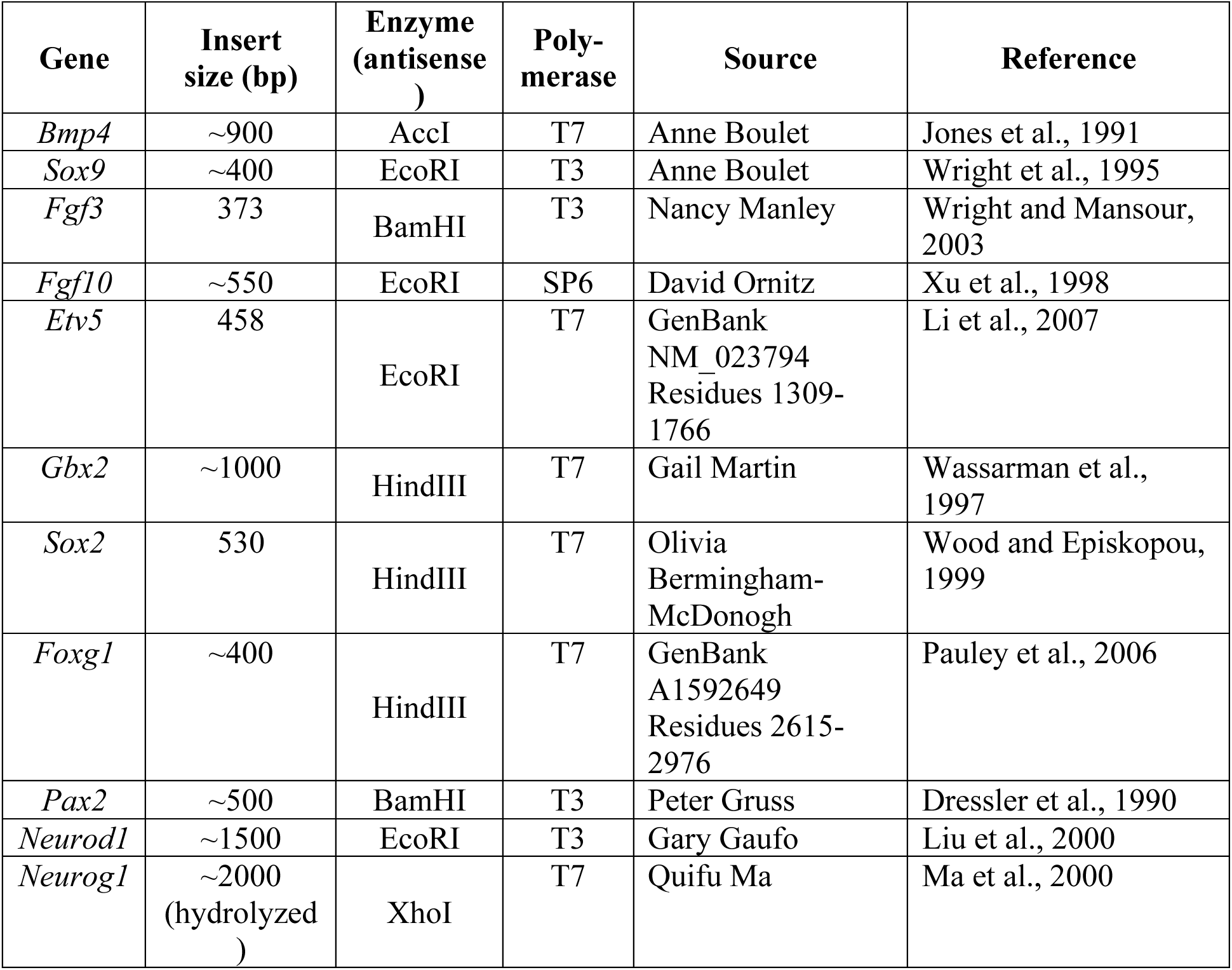
cDNA clones used to prepare digoxigenin-labeled cRNA antisense transcripts for in situ hybridization.

Purified plasmid DNA was digested with the indicated restriction enzyme and then transcribed with the indicated RNA polymerase to produce antisense probes for ISH.

**Table S2:**
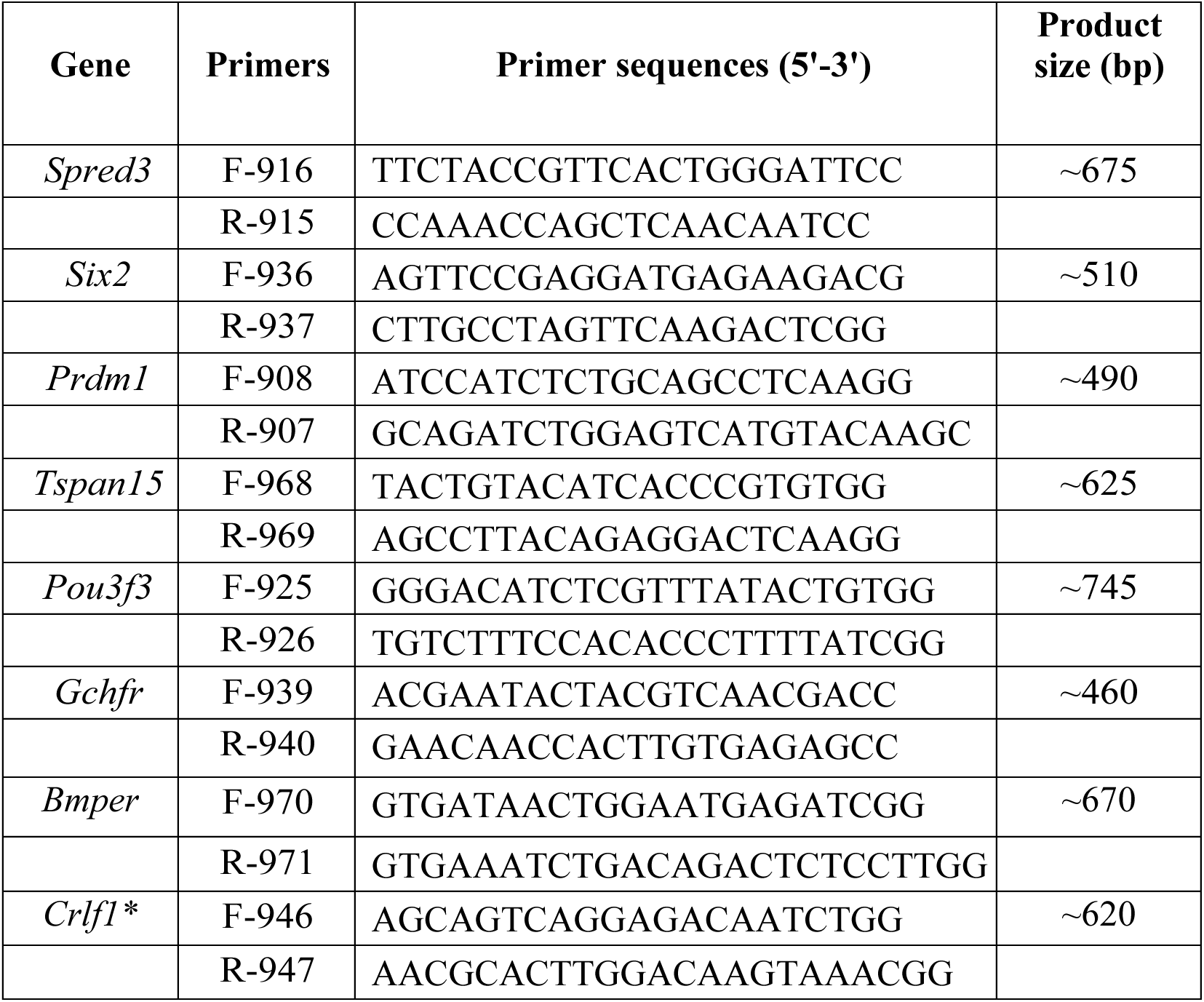
Primers used to generate DNA fragments for cRNA probe generation.

Forward (F) and reverse (R) primers used to PCR-amplify 3’UTR regions of each indicated gene from mouse genomic DNA, except *Crlf1*, which was amplified from a cDNA clone. All reverse primers include the T7 promoter sequence (GGATCCTAATACGACTCACTATAGGGAG) at the 5’ end. The antisense-RNA strand was produced by transcription of the PCR product using T7 RNA polymerase. *Crlf1* cDNA 3 from DNASU Arizona State University Clone # MmCD00295268 (NCBI NM_018827.2, Image: 100063851).

**Figure S1.**
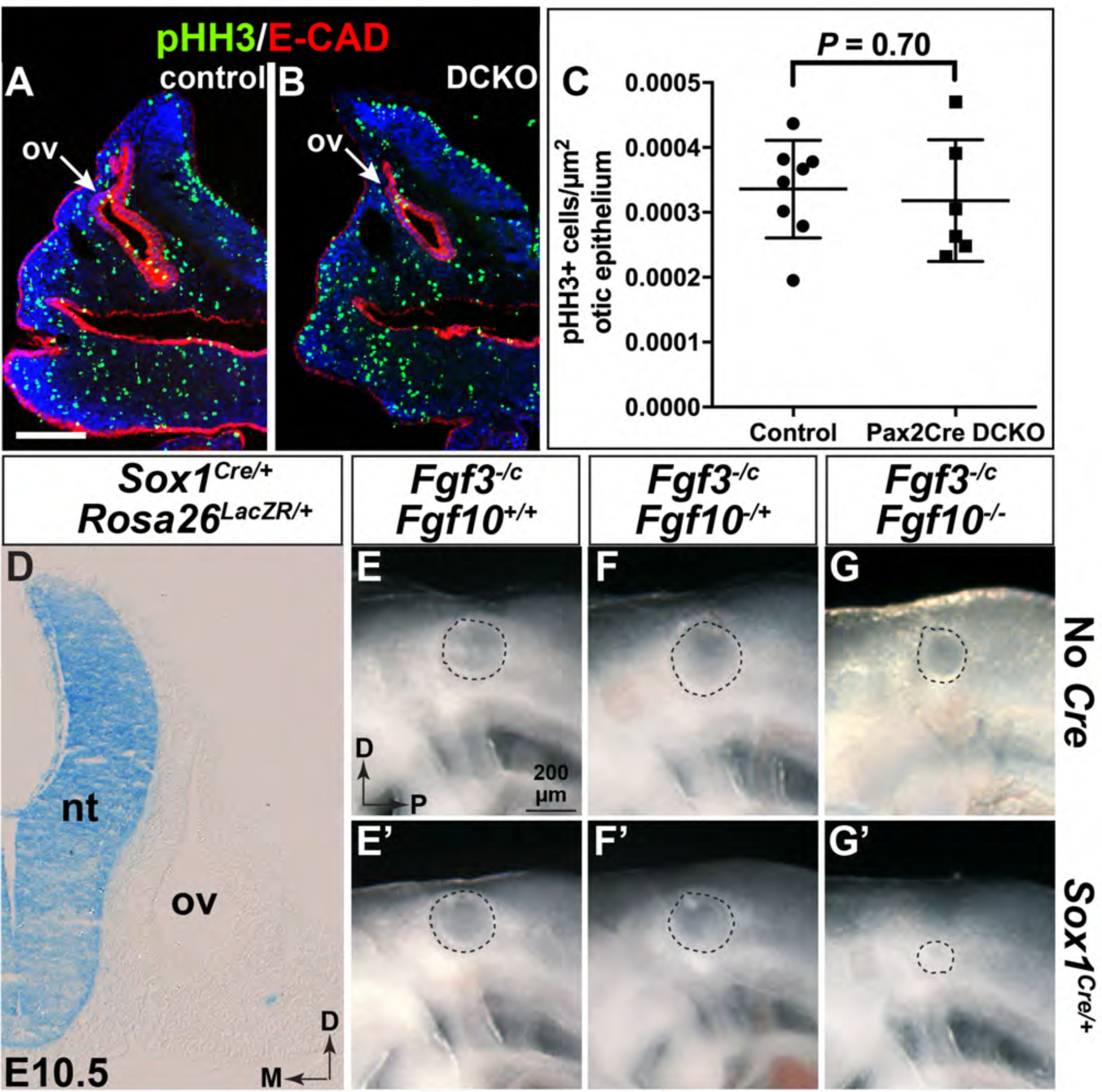
*Fgf3* and *Fgf10* are not required in the *Pax2-Cre* lineage for early otocyst proliferation. (A,B) Transverse cryosections of E10.5 control *(Fgf3^-/c^;Fgf10^-/c^)* and double conditional mutant (DCKO; *Fgf3^-/c^;Fgf10^-/c^;Tg(Pax2-Cre)/+)* otic vesicles (ov) immunostained to detect pHH3 (green) and E-Cadherin (red). (C) Quantification of pHH3-positive cells per otic epithelial area shows no significant difference between genotypes. N=8 control and 6 DCKO otocysts. Scale bar in A (100 μm) applies to B. (D) Transverse section at the level of the otic vesicle of an E10.5 X-gal-stained *Rosa26^LacZR/+^;Sox1^Cre/+^* embryo shows CRE activity restricted to the neural tube. (E-G’) Lateral views of otic vesicles from freshly dissected E9.5 embryos show that in the global absence of *Fgf10*, hindbrain *Fgf3* is required to develop a normally sized otic vesicle (G’). *Fgf* genotypes are indicated above and *Cre* status is to the right of each row. Dashed lines denote the external circumference of the otic epithelium. The scale bar and orientation axes in E apply to E-G’. Abbreviations: D, dorsal; M, medial, nt, neural tube; ov, otic vesicle; P, posterior.

**Figure S2.**
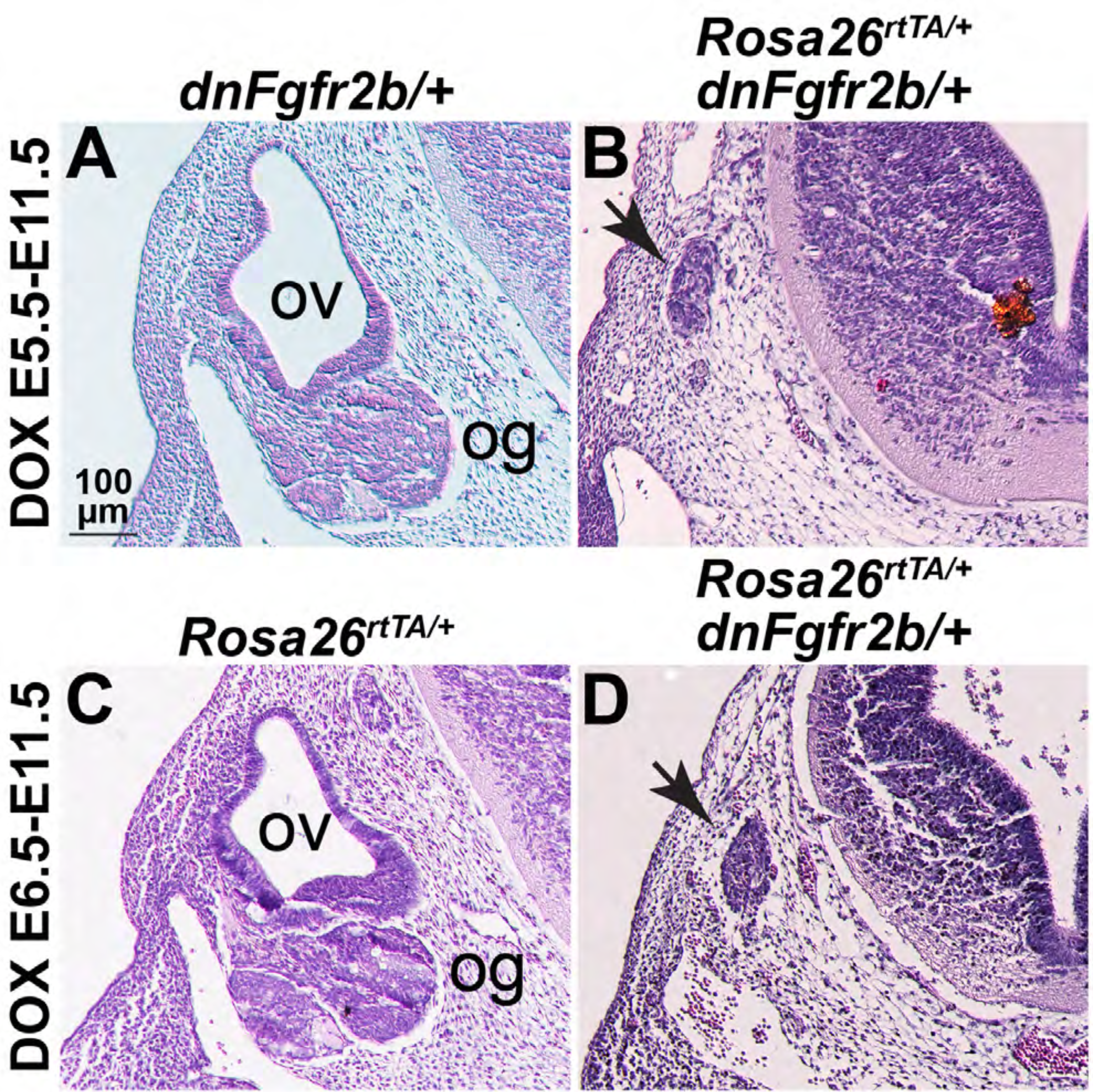
Induction of dnFGFR2b prior to otic placode induction blocks otic vesicle formation. Transverse hematoxylin and eosin-stained sections of embryos derived from Tg(tetO-*dnFgfr2b)/+* females crossed to *Rosa26^rtTA/+^* males and fed DOX-chow between E5.5-E11.5 (A,B) or E6.5-E11.5 (C,D). Genotypes are indicated above each column and DOX induction conditions indicated to the left of each row. Remnant otic tissue in the double heterozygotes is indicated with an arrow (B,D). The scale bar in A applies to all panels. Abbreviations: og, otic ganglion; ov, otic vesicle.

**Figure S3.**
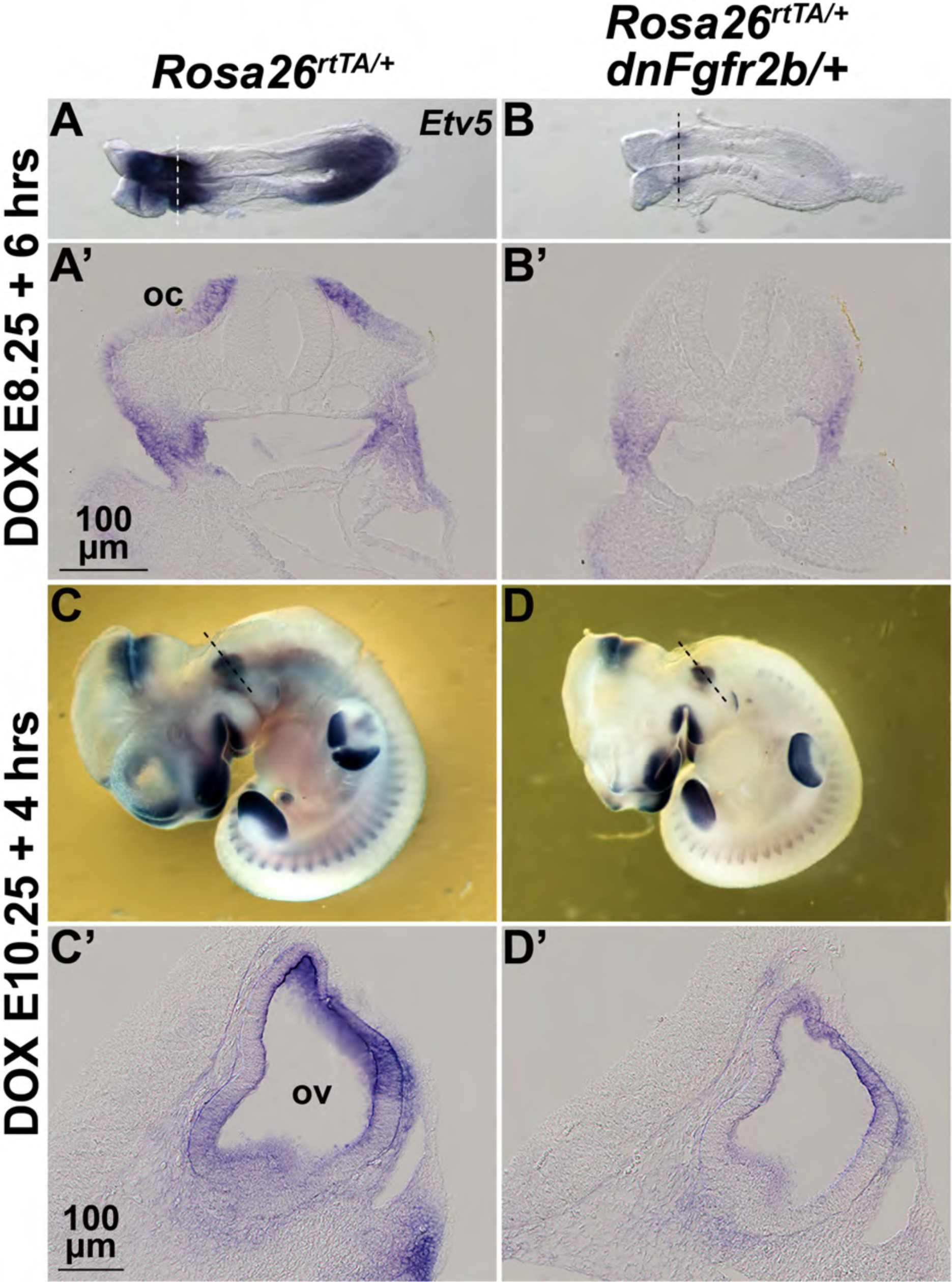
Inhibition of FGFR2b ligands has a rapid onset. Whole-mount ISH with an *Etv5* probe on embryos derived from wild type females crossed to *Rosa26^rtTA/rtTA^;Tg(tetO-dnFgfr2b)/+* males and treated with DOX for six hours beginning at E8.25 (A-B’) or 4 hours beginning at E10.5 (C-D’). Genotypes are indicated above each column. Dashed lines (A,B,C,D) indicate the planes of transverse sections shown in A’,B’,C’,D’. Abbreviations: oc, otic cup; ov, otic vesicle.

**Figure S4.**
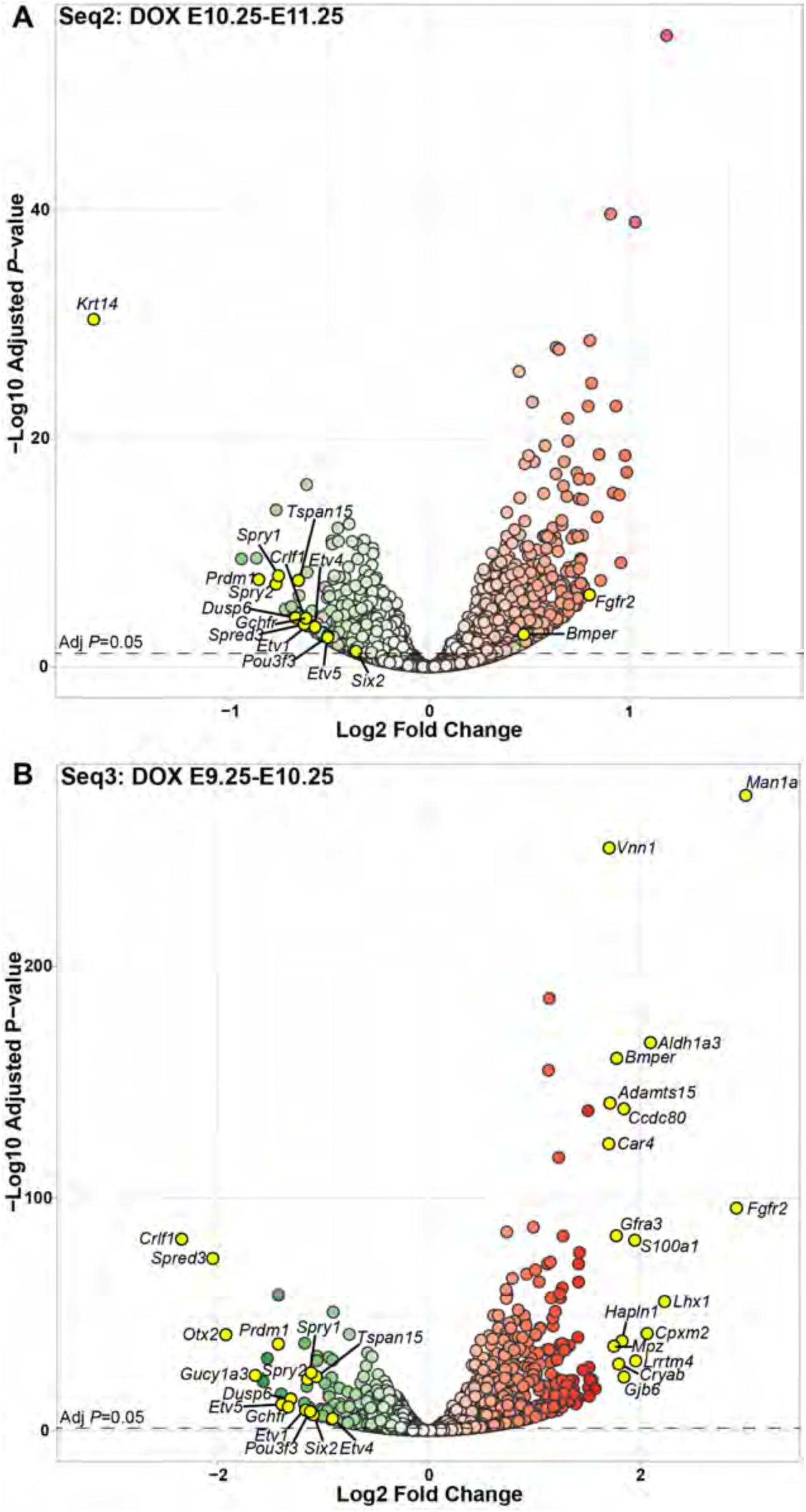
Differential RNA-Seq analysis of otic epithelia subjected to overlapping and longer dnFGFR2b induction windows define a stage-specific FGFR2b signaling response during early otic morphogenesis. (A) Volcano plot of the RNA-Seq2 (DOX exposure E10.25-E11.25), and (B) RNA-Seq3 (DOX exposure from E9.25-E10.25) datasets. Down-regulated (green) and up-regulated (red) genes were identified using a paired statistical model (see Methods). The statistical significance of the differential expression is shown on the y-axis and the fold change is shown on the x-axis. Names for genes highlighted in yellow include all those with a fold-change > 1.5, plus known FGF target genes and genes that we pursued for expression validation.

**Figure S5.**
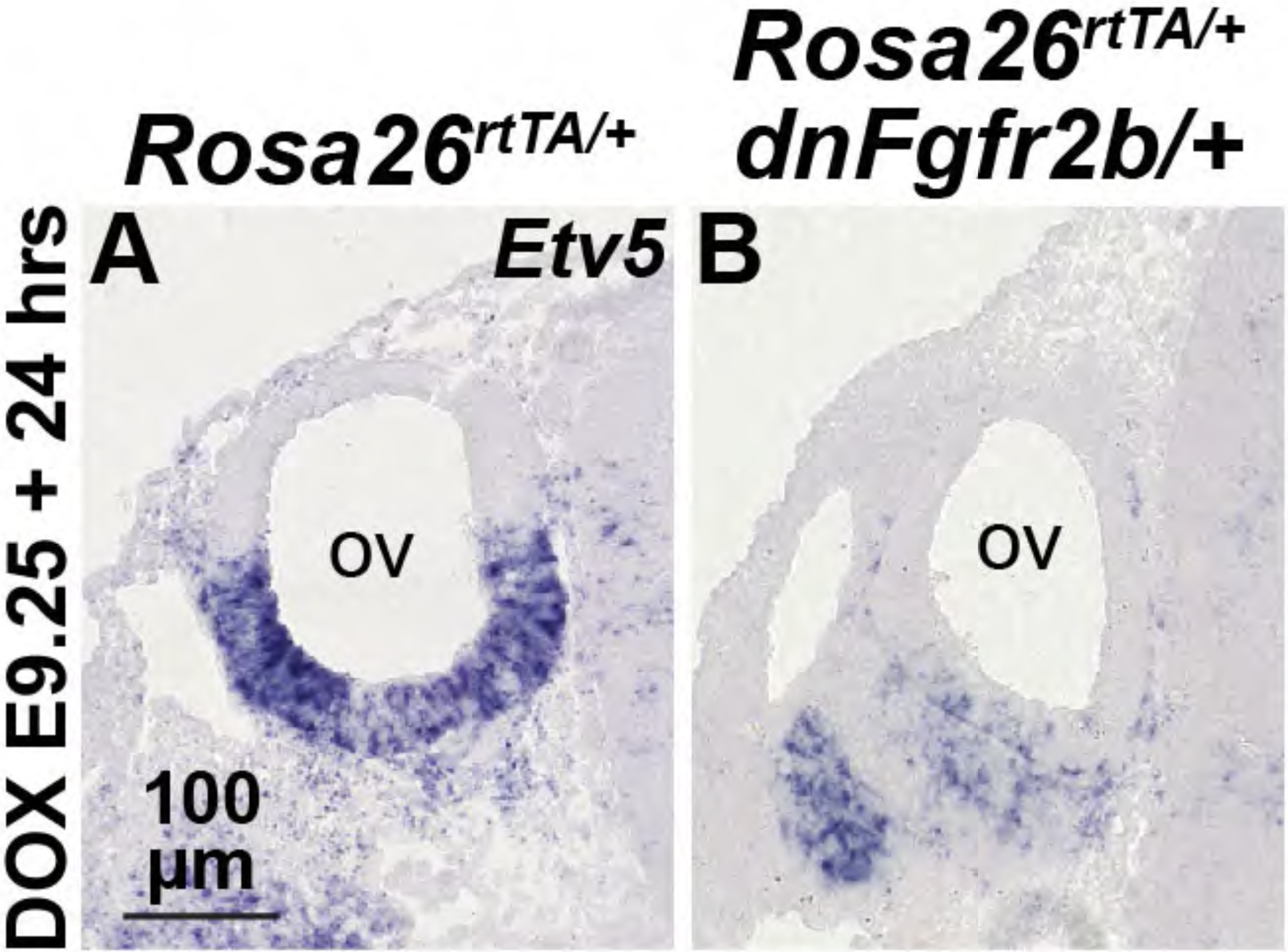
*Etv5* is lost from the otic vesicle following RNA-Seq3 dnFGFR2b induction conditions. ISH using an *Etv5* probe on sections taken through the otic vesicle (ov) and ganglion (og) from control (A) *(Rosa26^rtTA/+^)* or *dnFgfr2b-expressing* embryos (B) subjected to Seq3 induction conditions (E9.25-E10.25, n=3 each).

**Figure S6.**
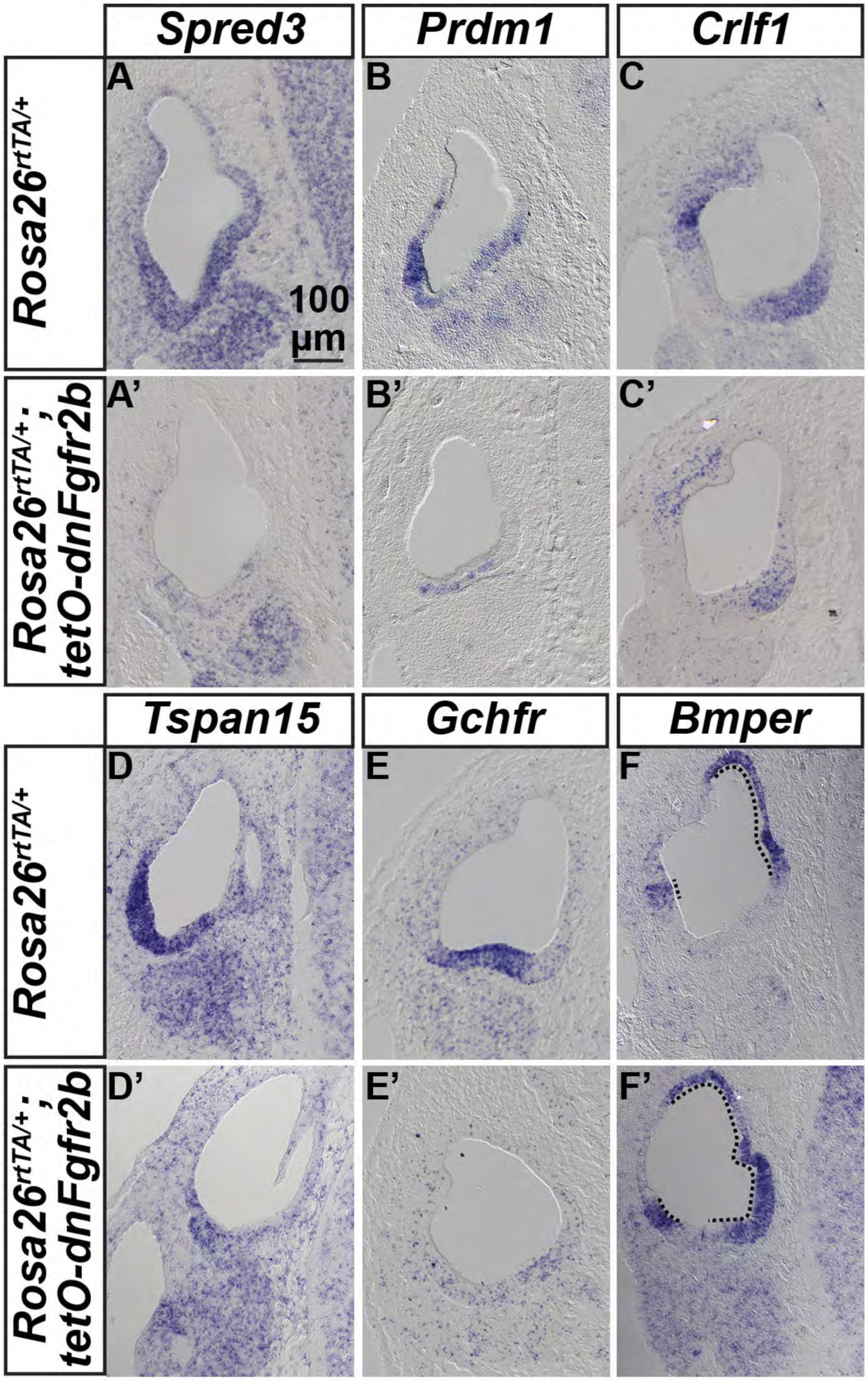
Validation of the subset of new FGFR2b ligand-dependent genes differentially expressed under RNA-Seq2 induction conditions. ISH of transverse E11.25 paraffin sections of otocysts exposed to Seq2 induction conditions (DOX, E10.25-E11.25). Probes are indicated in the boxes above each column and genotypes are indicated to the left of each row (controls A-F, dnFGFR2b-expressing embryos A’-F’, n=3 each). Dorsal is up, medial is to the right. The scale bar in A applies to all panels. The dotted line in F and F’ demarks *Bmper* expression.

**Figure S7.**
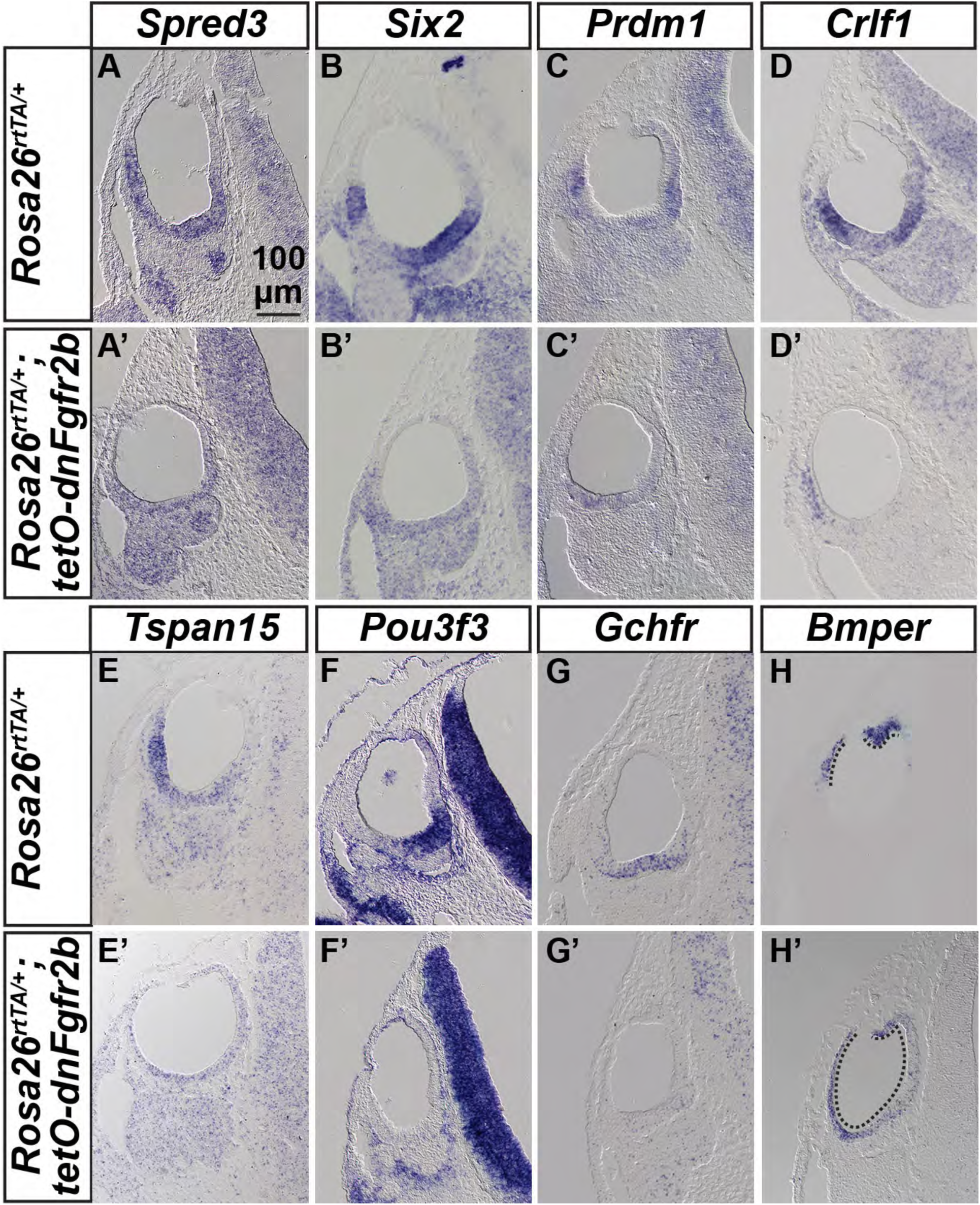
Validation of the subset of new FGFR2b ligand-dependent genes differentially expressed under RNA-Seq3 induction conditions. ISH of transverse E10.25 paraffin sections of otocysts exposed to Seq3 induction conditions (DOX, E9.25-E10.25). Probes are indicated in the boxes above each column and genotypes are indicated to the left of each row (controls A-H, dnFGFR2b-expressing embryos A’-H’, n=3 each). Dorsal is up, medial is to the right. The scale bar in A applies to all panels. The dotted line in H and H’ demarks *Bmper* expression.

